# Disruption of small RNAs and mechanistic variation in *Segregation Distorter—*a sperm-killing drive system in *Drosophila melanogaster*

**DOI:** 10.64898/2025.12.01.691737

**Authors:** Logan T. Edvalson, Xiaolu Wei, Ching-Ho Chang, Amanda M. Larracuente

## Abstract

Meiotic drivers achieve biased transmission to the next generation, often at the expense of their host. Drive is widespread and can shape the evolution of proteins, chromosome structure, and karyotype. The sperm killer *Segregation Distorter* (*SD*) in *Drosophila melanogaster* is a well-studied driver but like most complex drivers its mechanism remains elusive. *SD* is a multigene complex, frequently associated with chromosomal inversions, where the main driver locus, a truncated duplication of the gene *RanGAP* kills wild-type sperm containing a satellite DNA called *Responder* (*Rsp*). Functional small RNAs are frequently implicated in the mechanisms of sperm killers, and we recently showed that *Rsp* is a source of these RNAs. Here we use transcriptomics in two *SD* haplotypes to link *Rsp* expression and/or RNAs to drive. We found that *Rsp*-derived small RNAs are underrepresented in driving testes of only one of the *SD* haplotypes. We show that over-expressing *Rsp* is sufficient to reduce drive strength in the haplotype with downregulated *Rsp* but not the other. We therefore shed light on the mechanism of *SD* by making a connection between the target and the drive phenotype. Additionally, our data imply that different haplotypes of complex drivers, like *SD*, can vary in their mechanism.

## Introduction

Meiosis facilitates key principles of inheritance, including the fair segregation of alleles to gametes. However, selfish genetic elements, called meiotic drivers (1), can subvert fair segregation and bias their transmission to the next generation. Although some target meiosis in the strict sense, meiotic drivers can act through a diversity of mechanisms during any stage of gametogenesis (2). Meiotic drive is widespread across species and can have major consequences for genome evolution, gametogenesis, and even speciation (reviewed in 3–5). Meiotic drivers can impose fitness costs on their hosts directly as a consequence of drive (e.g. fertility costs) or indirectly through linked deleterious mutations (e.g. 6–9). The costs of drive apply selective pressure on the host to evolve suppressors that restore fair segregation, which then in turn can prompt drivers to evolve ways to circumvent host suppression (4). This tit for tat dynamic can result in perennial arms races between meiotic drivers and host genomes, triggering rapid evolution and innovation of sequences involved in the conflict. As a consequence, drive systems can be complex, involving enhancers and suppressors across the genome (10). Male systems often involve similar phenotypes (*e.g.* sperm chromatin) and frequently target repeat-rich heterochromatin, suggesting that they may exploit common vulnerabilities in spermatogenesis (11).

One emerging theme across meiotic drive systems is the involvement of RNA interference (RNAi) pathways that mediate gene and repeat silencing across organisms (11). RNAi is implicated in several drive systems on both sides of the genetic conflict. For example, small interfering RNAs with sequence complementarity to drivers can act as drive suppressors: Sex ratio distorters like *Stellate* in *D. melanogaster*, and the *Winters* and *Durham* systems in *D. simulans,* involve loci that produce either piwi-interacting RNAs (piRNAs) (12) or hairpin RNAs (hpRNAs) (13) that act as drive suppressors. Similarly, the *kinesin driver (Kindr)* complex in maize promotes the biased segregation of heterochromatic knobs and is suppressed by small interfering RNAs (siRNAs) (14). RNAi pathways are also implicated beyond drive suppression. For example, in the Paris sex ratio system of *D. simulans,* driving X chromosomes disrupt Y chromosome segregation. One of the essential drive loci in this system is *HP1D2*, a paralog of *Rhino*—a gene important for producing piRNAs (15). More broadly, duplications of RNAi-related genes appears common on sex chromosomes across taxa (16,17), suggesting that dose-sensitive genetic conflicts involving RNAi may be widespread and play important roles in sex chromosome evolution (18,19).

While some common themes are emerging from studies across meiotic drive systems (reviewed in 11), we currently know little about precise molecular mechanisms of drive. Many drive systems are complex—involving multiple loci, gene duplications, and repetitive DNA—and are often associated with regions of suppressed recombination (19) making it difficult to identify and study how key components of drive systems interact.

Here we use a classical system that shares common features with other male drivers (e.g. defects in sperm chromatin, gene duplication, and repetitive DNA) to gain new insights into drive mechanisms: the *Segregation Distorter (SD)* of *D. melanogaster*. We use *SD* to study molecular features of male meiotic drive at the RNA level*. SD* is a complex autosomal sperm-killing driver segregating on the second chromosome of populations across the globe (20). *SD* biases its transmission by killing sperm bearing sensitive alleles of its target, a repetitive array of tandem satellite DNA repeats in the pericentromeric heterochromatin called *Responder (Rsp)* (21,22). The main drive locus corresponds to a partial tandem duplication of *Ran GTPase Activating Protein* (*Sd-RanGAP*; 23,24). *RanGAP* is a GTPase activating protein involved in the *Ran* cycle, which is important for multiple cellular processes, including nuclear transport, spindle assembly, and nuclear envelope assembly (25). *SD* is a multi-gene complex: all drive haplotypes carry *Sd-RanGAP,* an insensitive allele of the *Rsp* target locus, and multiple enhancers that strengthen drive across different *SD* haplotypes, e.g. *Enhancer of SD (E(SD))* (26), *Modifier of SD (M(SD))* (27), and *Stabilizer of SD (St(SD))* (28). We do not yet know the molecular identity of any drive modifiers. To enhance linkage between *SD* and their enhancers, most *SD* chromosomes acquire specific chromosomal inversions to suppress recombination (reviewed in Larracuente and Presgraves 2012). Therefore, despite sharing the *Sd-RanGAP* locus, *SD* chromosomes may differ in their inversions and modifiers, and thus show different levels of distortion and cytological defects (30).

The classic *SD* drive phenotype manifests as a post-meiotic chromatin defect (31,32), but the mechanism is unknown. The driver, *Sd-RanGAP*, is enzymatically active, but mislocalized to the nucleus, which likely disrupts *Ran*-mediated nuclear transport (23). In contrast, we know little about the role of the target in drive. *Rsp* is a pericentromeric satellite DNA array on chromosome 2R consisting of tandemly repeated dimers of two related ∼120-bp repeats: *L-Rsp* and *R-Rsp* (21,22). The sensitivity of a chromosome to drive correlates positively with the copy number of *Rsp* (21,22,33). The drive sensitivity of 2^nd^ chromosomes range from insensitive (<100 copies of *Rsp*), to sensitive (∼300 to ∼1500 copies), or super sensitive (> ∼2000 copies) (22). Yet, we do not know what features of *Rsp* make it a target of drive. Satellite DNAs may also be targets of other drive systems. For example, many sex ratio distorters target Y chromosomes, which are rich in satellite repeats that vary in copy number and composition (reviewed in 11). A deeper understanding of the roles that satellite DNAs play in drive may reveal a common vulnerability in spermatogenesis, one that selfish genetic elements repeatedly exploit.

While previously regarded as inert components of the genome, complex satellite DNAs like *Rsp* are transcribed during gametogenesis by specialized heterochromatin-dependent transcriptional machinery (34,35). Transcripts from *Rsp* are primarily processed into small 23-30 bp piRNAs in both ovaries and testes and may contribute to heterochromatin formation in the early embryo (34). Others have identified potential functions for other satellite DNA-derived RNAs in gametogenesis (*AAGAG*, 36,37) and X chromosome recognition in dosage compensation (*1.C88*, 38) but we know little about what functions, if any, *Rsp-*derived piRNAs may have during gametogenesis, particularly spermatogenesis.

Several lines of evidence set the precedent that *Rsp*-derived RNAs may have a role in drive. First, piRNAs are involved in chromatin remodeling in the germline (39), and the *SD* phenotype involves a chromatin defect (30–32). Second, mutations in the piRNA pathway component *Aubergine* enhance *SD* drive (40). Finally, non-coding RNAs originating from pericentromeric satellite DNAs can play a role in sperm development (36,37). Knocking down (36) or disrupting the transcription (37) of an abundant simple satellite, *AAGAG,* affects a post-meiotic sperm chromatin and morphological defect (37), similar to the defect in *SD-*targeted sperm (30–32). While the role of satellite DNA-derived RNAs is still unclear, they may contribute to chromatin regulation during spermatogenesis. We propose that meiotic drivers like *SD* may exploit these features of satellite DNA to bias their transmission.

Here we test the hypothesis that *Rsp*-derived RNAs are involved in drive by exploring the total and small RNA transcriptomes of driving testes with two different *SD* haplotypes that differ in their chromosomal inversions (*SD-Mad* and *SD-5*). We discovered a significant dearth of *Rsp-*derived small RNAs in driving genetic backgrounds associated with one drive haplotype (*SD-Mad*) but not another (*SD-5*). To confirm that *Rsp* small RNAs are involved in *SD-Mad* drive, we overexpress *Rsp* and show corresponding changes in drive strength. Our results are consistent with a haplotype-specific role for *Rsp*-derived RNAs in drive and suggest a potential role for complex satellite-derived RNA in spermatogenesis. We propose that different drive haplotypes may vary in their modifiers, giving rise to molecular variation in drive.

## Results

### Rsp-derived small RNAs are underrepresented in the small RNA transcriptome of SD-Mad testes

Because there is evidence that satellite-derived small RNAs (smRNAs) are important for sperm development (36,37), we hypothesized that *SD* may interact with these smRNAs as a part of its mechanism. We explored the small RNA transcriptome of testes heterozygous for combinations of two different driving *SD* haplotypes (*SD-Mad* and *SD-5*) (Figure 1A) and wild type chromosomes with different *Rsp* alleles (Figure 1A). Because our previous work found a correlation between *Rsp* copy number and RNA abundance (34), all of our comparisons are relative to *Rsp* copy number controls (*R16/Gla* or *R16/Iso1* where *R16* contains a *Rsp* deletion allele and *Iso1* and *Gla* contain a sensitive and super-sensitive *Rsp* allele, respectively; Figure 1A) (26). The *SD* haplotypes we use differ by their inversion types (Figure 1A) but both exhibit strong drive, measured as *k,* or the proportion of *SD* offspring from a cross with *SD* heterozygous males (Table S1).

**Figure 1:**
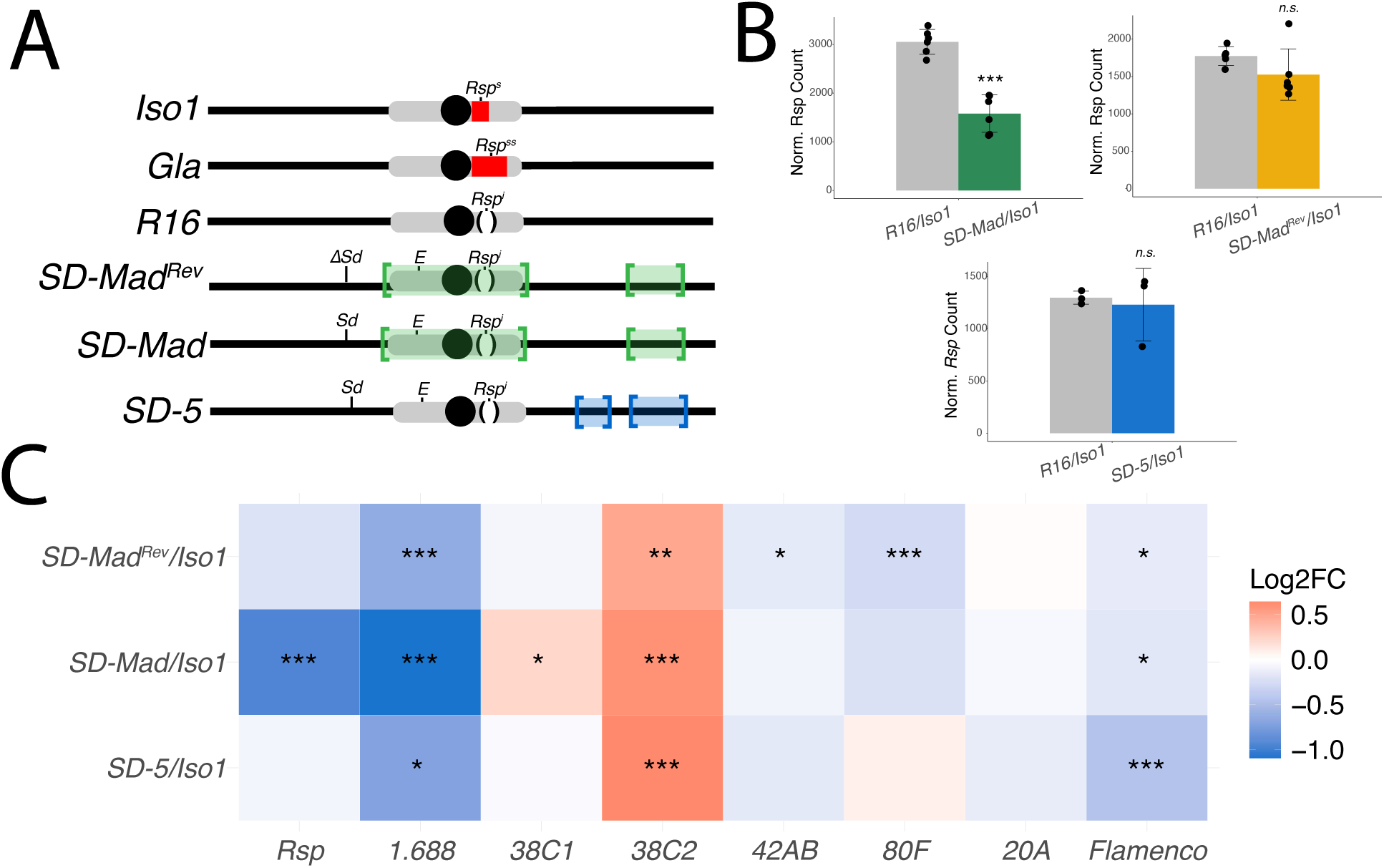
Analysis of small RNAs in SD-Mad heterozygotes show a lower abundance of small RNAs corresponding to Rsp. A) A schematic of the 2nd chromosomes used in this study. Iso1 and Gla are sensitive backgrounds with varying copy number of Rsp. Iso1 contains ∼1100 copies of Rsp making it sensitive and Gla contains >3000 making it super sensitive. R16 is a copy number control for SD and does not drive. SD-Mad and SD-5 are driving chromosomes containing different inversions (highlighted in colored boxes). They contain the main drive locus, Sd-RanGAP and show perfect or near perfect drive against sensitive chromosomes. SD-MadRev is a haplotype control for SD-Mad. It is identical to SD-Mad with the exception of a deletion of Sd-RanGAP. B) A comparison of the DESeq2 normalized counts of Rsp in each comparison shows that Rsp smRNAs are less abundant in SD-Mad heterozygotes relative to R16 heterozygotes (Log2FC = -0.95, P.adj = 5.1e−9). Rsp smRNAs are apparently unaffected in SD-MadRev heterozygotes indicating that the reduction of smRNAs in SD-Mad heterozygotes is correlated with drive. Rsp RNAs are also unaffected in the SD-5 background indicating that SD-5 does not perturb Rsp smRNA abundance. C) A heatmap showing the differential abundance of smRNAs for major piRNA source loci in the genome. 1.688, flamenco, and 38C2 are differentially expressed in this background.

We found significantly fewer *Rsp-*derived smRNAs (piRNAs and endo-siRNAs) in some driving testes. *Rsp*-derived smRNAs are significantly less abundant in the testes of *SD-Mad/Iso1* heterozygotes relative to the non-driving copy number control (*R16/Iso1*) (Figure 1B; Log2FC = -0.95, *P.adj* = 5.1e^-9^, DESeq2). To confirm that the low abundance of *Rsp* smRNAs correlates with drive rather than non-drive related factors in the genetic background, we created a revertant *SD-Mad* background (*SD-Mad^Revertant^* or *SD-Mad^Rev^*, Figure 1A). We used CRISPR/*Cas9* to delete the *Sd-RanGAP* duplication from *SD-Mad*, leaving one full length copy of RanGAP and the rest of the chromosome with putative drive modifiers intact. This chromosome is therefore identical to *SD-Mad* outside of *Sd-RanGAP* but no longer exhibits strong drive (*k* =* 0.54). We expect that any differences in smRNA abundance between *SD-Mad/Iso1* and *SD-Mad^Rev^/Iso1* are therefore due to drive or *Sd-RanGAP* itself. Indeed, we found that *Rsp* smRNAs are more abundant in *SD-Mad^Rev^/Iso1* relative to *SD-Mad/Iso1* (Log2FC = 0.66, *P.adj* = 0.004, DESeq2) and at similar levels as the copy number control, *R16/Iso1* (Figure 1; Log2FC = -0.21, *P.adj* = 0.19, DESeq2). This indicates that the difference in *Rsp* smRNA abundance in *SD-Mad/Iso1* is likely due to the drive phenotype. We see a similar dearth of *Rsp-*derived smRNA in driving testes in an independent background with a super-sensitive *Rsp* allele (*SD-Mad/Gla)* compared to its non-driving copy number control (*R16/Gla)* (Figure S1; Log2FC = -0.68, *P.adj* = 2.6e^-14^, DESeq2). This suggests that the effects on *Rsp* smRNA abundance are related to the driving *SD-Mad* chromosome rather than the target chromosome.

To determine if the effect on *Rsp* smRNAs is specific to *SD-Mad,* we explored the small RNA transcriptome in driving testes with another *SD* haplotype (*SD-5*) that differs in its chromosomal inversions (29). Intriguingly, *Rsp* smRNA abundance is similar between *SD-5* and *R16* in an *Iso1* background (Figure 1; Log2FC = -0.07*, P.adj* = 0.79, DESeq2) and higher than expected levels in the *Gla* background (Figure S1; Log2FC = 0.33, *P.adj* = 2.3e^-11^, DESeq2). Therefore, the disruption of *Rsp* RNAs is not a universal feature of *SD* chromosomes.

Outside of *Rsp,* we see shifts in many repeat-derived smRNAs in driving testes of both *SD* haplotypes: for example, 30% (12/40) and 27.5% (11/40) of repeat annotations (satellites DNAs, piRNA clusters, and TEs) were differentially expressed (*P.adj* ≤ 0.01) in *SD-Mad* and *SD-5*, respectively (Figure S2; with similar—56.4% (22/39) and 50% (20/40)—effects in the *Gla* background; supplementary file 1). There are only a few notable patterns that arise from these comparisons (Table S2). We found that another complex satellite DNA on the X chromosome, presumably unrelated to *Rsp*, *1.688*, is significantly less abundant in all *SD* heterozygote testes in the *Iso1* background comparisons (Figure 1C, Table S2; Log2FC*^SD-Mad^* = -1.1, *P.adj^SD-Mad^* = 4.8e^-14^; Log2FC*^SD-MadRev^* = -0.64, *P.adj^SD-MadRev^* = 1.16e^-12^; Log2FC*^SD-5^* = -0.71, *P.adj^SD-5^* = 0.02, DESeq2) and in *SD-Mad/Gla* testes (Figure S1; Log2FC*^SD-Mad^* = -0.28, *P.adj^SD-Mad^* = 0.001). SmRNAs derived from a piRNA cluster linked to *Sd-RanGAP* on chromosome 2L, *38C2*, and the X-linked somatic piRNA cluster, *ffamenco,* is differentially expressed for all genotypes in both backgrounds except for *Flamenco* in *SDMad/Gla* (Figure 1C, S1). There is no evidence that either *38C2* or *Flamenco* are involved in *SD-*mediated drive.

Our results demonstrate that *SD-Mad* and *SD-5* haplotypes, despite sharing the same main drive locus, have different effects on smRNAs derived from repetitive loci such as complex satellites (including *Rsp*), transposable elements, and piRNA clusters. Since these *SD* chromosomes differ in their inversions, we propose other linked loci might affect the drive mechanisms and phenotypes.

The dearth of *Rsp* smRNAs in *SD-Mad* heterozygotes could be due to a disruption in transcription of the locus or subsequent processing steps. Many factors can influence piRNA production. For example, the piRNA pathway can amplify piRNAs independently of transcription, such as the ping pong cycle, (41). Notably, *Rsp* piRNAs do not have a strong ping pong signature in testes (34,42). To distinguish between a disruption in transcription or some downstream process, we examined total RNA.

### Broad differences in the long RNA transcriptome of different driving backgrounds

We performed total RNAseq to determine how the expression landscape of coding and non-coding sequences differs in testes with drive. We found that 24.2% (3359/13825) of the total RNA transcriptome is differentially expressed (*P*.adj ≤ 0.01) in the *SD-Mad* background (*SD-Mad/Gla* vs *R16/Gla*; Figure S3B). Driving testes with *SD-5* had similarly divergent transcriptomes compared to copy number controls (24.3% (3352/13747) DEG in *SD-5/Gla* vs R16/Gla; Figure S3E). Repetitive loci were more frequently differentially expressed (*P*.adj ≤ 0.01) in *SD-Mad* heterozygotes compared to *SD-5* heterozygotes (41% (16/39) vs 23% (8/40), Figure S4, Table S3). Despite the effect on *Rsp* smRNA abundance in *SD-Mad* heterozygotes, the abundance of *Rsp* long non-coding RNAs (lncRNAs) does not show any consistent pattern between the backgrounds (Figure S5A-B, Table S3). This could be due to low counts for *Rsp.* On the other hand, we did find some differences in repetitive elements related to rDNA (R1, R2, and IGS) and *Tc1-Mariner* family TEs (all backgrounds; Figure S6). Interestingly, there was no correlation between the expression of TEs and the expression of piRNA clusters that contain fragments of these TEs in the total RNA, nor was there any correlation between the small RNAs from piRNA clusters and the total RNAs for those TEs. PiRNA clusters are usually defined in one isolate of *Iso1:* rapid turnover of TEs and piRNA sources could explain why we do not see a correlation between piRNA cluster expression and TE expression in our backgrounds.

The reduction in Rsp smRNAs without any corresponding change in the lncRNA in *SD-Mad*-mediated drive is intriguing, as piRNA precursors should be among these RNAs. Because previous studies based on immunofluorescence in testes showed that *Sd-RanGAP* is mislocalized in driving spermatocytes (23), we asked if the difference in smRNAs could be due to a mislocalization of the lncRNA. We performed RNA FISH for *Rsp* and the *1.C88* satellite as a fluorescence control in testes of *SD-Mad* heterozygotes. The most abundant variant of *1.C88* is the *35S-bp* satellite on the X chromosome, which is the same in our comparisons. Curiously, we do see some apparent differences in the pattern of 1.688 and, to a smaller extent, *Rsp* signals between driving and non-driving genotypes. However, the signals are nuclear (Figure S7), which suggests that mislocalization of the lncRNA, at least at a gross scale, may not explain the reduction in *Rsp* smRNAs.

We performed complementary analyses to identify patterns in gene expression. Genes make up 99.6 – 99.8% of the differentially expressed sequences in driving testes (Supplemental File 1). However, many of these differences likely reflect divergence among the second chromosomes used in each condition rather than the drive phenotype itself. To address this, we focused on pathway-level changes and coordinated gene expression rather than individual genes.

We conducted gene set enrichment analyses using PANGEA (43), and compared all differentially expressed genes (*P* ≤ 0.01) to Reactome gene sets (44) (Supplemental File 2). These analyses revealed a consistent enrichment in terms related to metabolism, the immune system, and the *Rho* GTPase cycle across driving testes (Figure S8). In addition, genes involved in sperm individualization (e.g. *asp, didum, bug22, chic, ctp, jar, nsr*) (45) were differentially expressed in both *SD-Mad/Gla* and *SD-5/Gla* (Figure 2A, Supplemental File 3), which is consistent with the hallmark chromatin defect (30–32).

**Figure 2:**
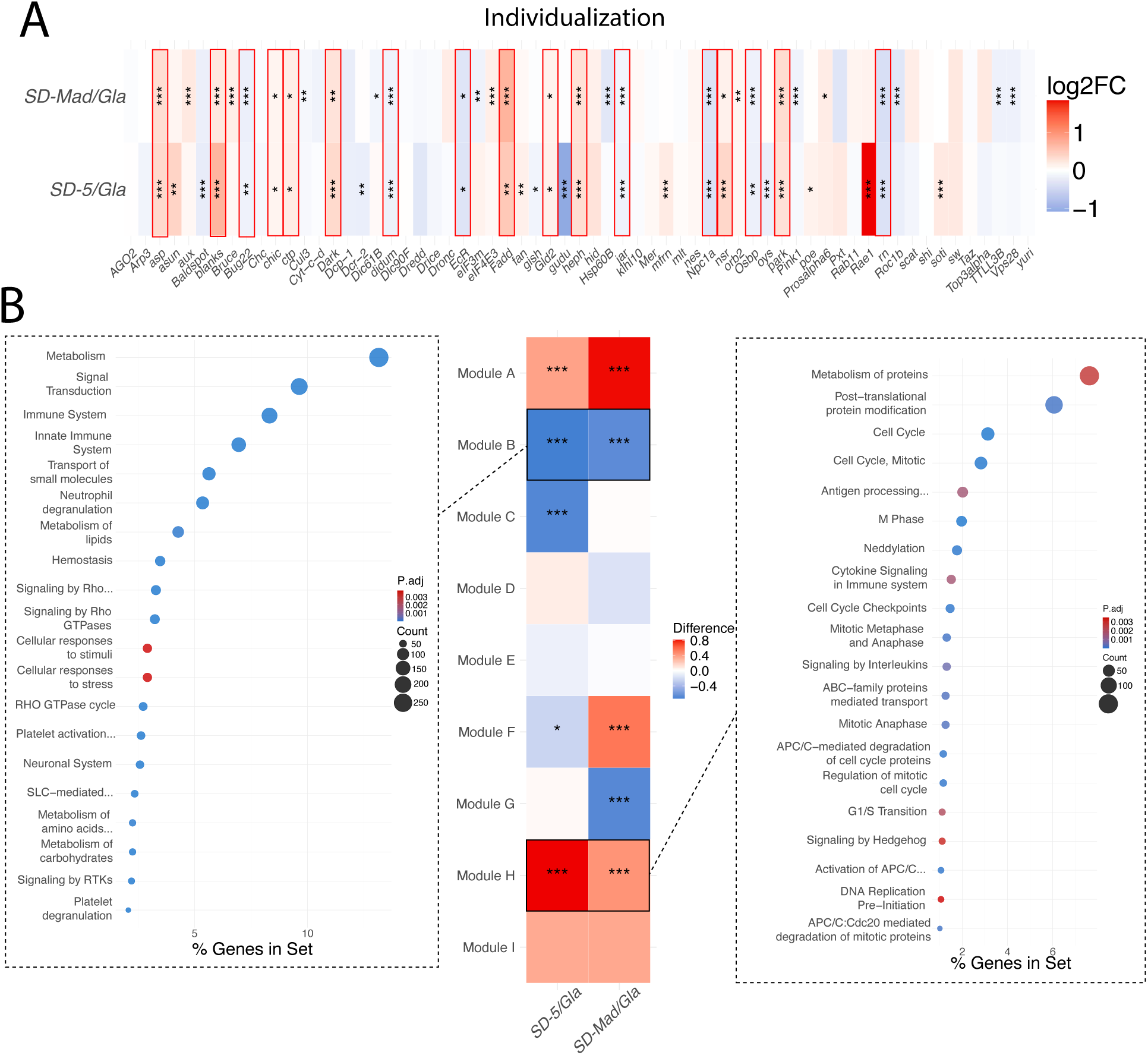
Analysis of total RNA from SD heterozygotes shows a broad dysregulation of genes involved in individualization and the cell cycle. A) A heatmap showing the direction and significance of differential expression for genes whose perturbation has been shown to result in defects at the individualization stage (Steinhauer 2015). Boxes outlined in red are genes that are differentially expressed in both SD haplotypes and in the same direction. B) WGCNA analysis of the total RNA identified modules of genes that are upregulated or downregulated in SD heterozygotes relative to the R16/Gla control. These modules were enriched for genes in Reactome gene sets. Module B contained genes that are downregulated and which are in gene sets related to metabolism, the immune system, and the Rho GTPase cycle. Module H contained genes that are upregulated and which are in gene sets related to the cell cycle and post translational modification.

To further identify coordinated expression patterns, we performed weighted gene co-expression network analysis (WGCNA) (46). WGCNA identified modules of co-expressed genes that distinguish driving genotypes from *R16/Gla* (Figure 2B, Table S4, Supplemental File 2). Gene set enrichment analysis of these modules using PANGEA (43) revealed that the drive phenotype is negatively correlated with several terms related to metabolism, the *Rho* GTPase cycle, and the immune system (Module B, Figure 2B).

Together, both pathway-level and network-based analyses implicate disruptions in cytoskeletal organization and/or sperm individualization. Whether these effects are directly related to drive mechanisms or downstream consequences remains unclear, but they are consistent with known phenotypes of spermiogenic failure.

WGCNA also identified two modules that were upregulated in driving genotypes. Our gene set enrichment analysis with PANGEA showed that one module (Module A) was not associated with any enriched pathways, whereas the other (Module H) was enriched for genes involved in the cell cycle and post-translational protein modification (Figure 2B).

*Rsp* is expressed well before individualization (34), during the period when male germ cells are actively dividing. If *Rsp* RNAs indeed contribute to the *SD* mechanism, the enrichment of cell cycle-related genes in Module H is consistent with a mechanism that originates earlier in spermatogenesis, around or before meiosis. Although the phenotype manifests after meiosis, temperature shift experiments suggest that the critical window for *SD* activity occurs before or during meiosis (47). Together, these results further support a model in which *SD* acts early in spermatogenesis, with defects only becoming apparent later, during individualization.

To identify genes that might interact to cause drive, we compared the gene expression of *SD-Mad/Iso1* to *SD-Mad^Rev^/Iso1.* These genotypes only differ by the presence of the main drive locus, *Sd-RanGAP. We* performed both totRNA and 3’ Digital Gene Expression (DGE) RNA sequencing and examined the overlap in differential expression between the totRNA and DGE sequencing. There are 69 differentially expressed genes where the DGE comparison is significant (*P^DGE^* ≤ *0.01*), and the sign of the Log2FC of the totRNA matches that of the DGE. Among this set of differentially expressed genes, 57 show at least a 50% difference in gene expression (absolute Log2FC value of at least 0.58 in DGE). These genes are not enriched in any Reactome gene sets. The top 20 most differentially expressed genes consists of 9 lncRNAs (3 anti-sense RNAs) and 11 protein coding genes: 8 of which are uncharacterized. The 3 characterized genes are *Artemis (Arts)*, *Gr61a,* and *Tono* (Figure S9, Supplemental File 1).

### *Loss of Kipferl results in the upregulation of satellite DNAs and weaker SD-Mad* drive

Our RNA seq analysis suggests that the reduction in *Rsp* smRNAs in *SD-Mad/Iso1* may be involved in the mechanism of drive. To test this hypothesis, we sought to determine if overexpressing *Rsp* is sufficient to rescue wild-type spermatids from *SD*. If drive is a consequence of reducing *Rsp* smRNAs in an *SD-Mad* background, then overexpressing *Rsp* RNAs in *SD-Mad/Iso1* testes may reduce drive strength. To overexpress *Rsp,* we took advantage of a recently-discovered component of the piRNA pathway called *Kipferl (kipf)* (48). In ovaries, *kipf* recruits piRNA transcriptional machinery to so-called *kipf-*dependent piRNA clusters. In its absence, the piRNA machinery becomes enriched at complex satellite DNAs, including *Rsp* and an unrelated repeat called *1.C88,* leading to their upregulation (48) (Figure 3A-B). While previous studies did not detect evidence for *kipf* expression in testes (49,50) (Figure S10), we found that *kipf* transcripts are indeed present in testes, albeit at low levels (Figure S10). We also found that, like in ovaries, satDNAs are overexpressed in *kipf* knock out (KO) testes, as demonstrated by reverse transcription quantitative polymerase chain reaction (RT-qPCR) (Figure 3B; testis fold change: *359-bp*, a component of the *1.C88* complex satellite = 2.85, *P* = 0.064; *Rsp* = 1.94, *P* = 0.038, t-test).

**Figure 3:**
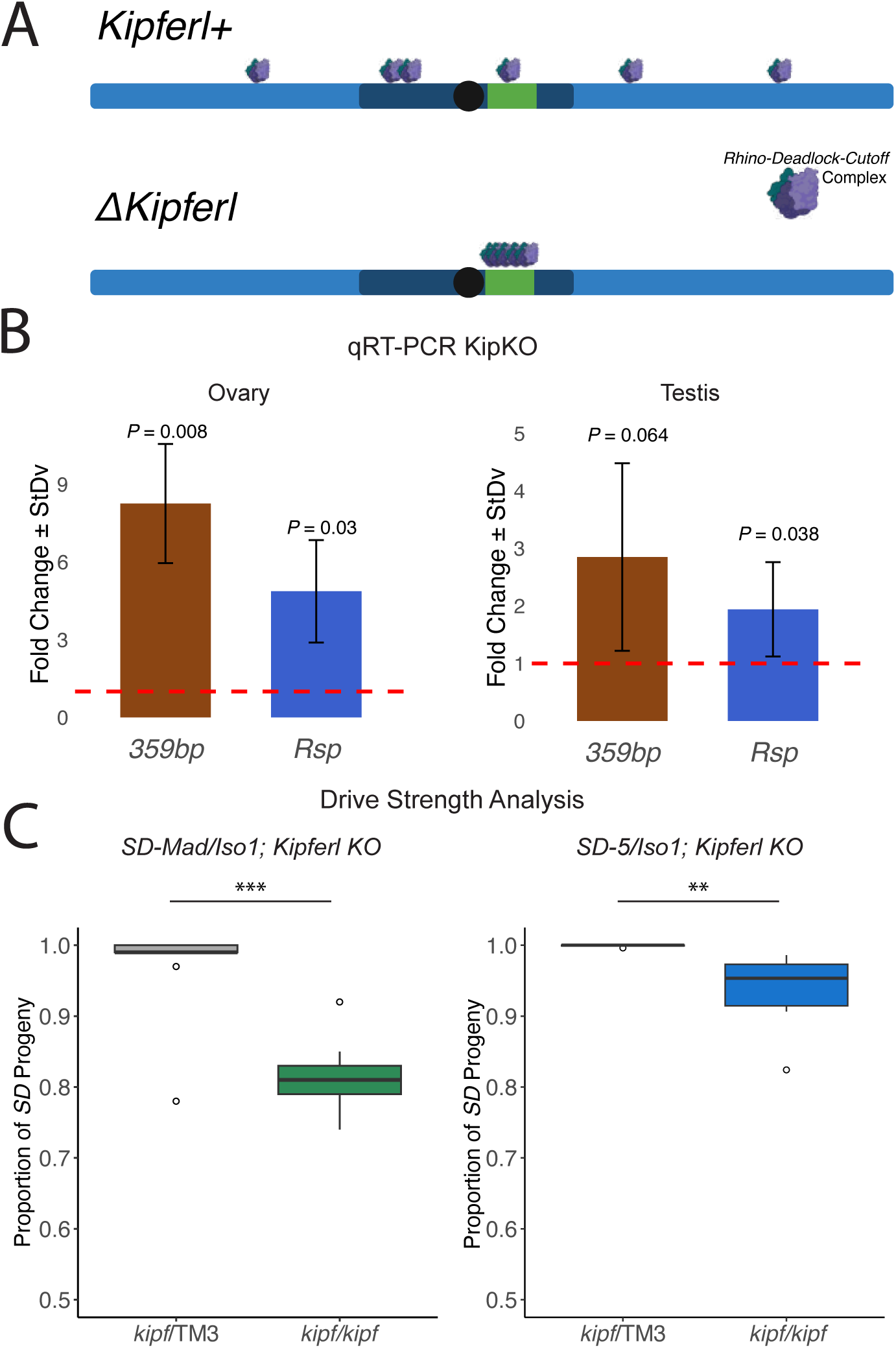
Satellite DNA overexpression via kipf-KO is sufficient to rescue drive in SD-Mad heterozygotes. A) A schematic of how we expect the localization of the RDC complex, the piRNA transcriptional machinery, to change upon knockout of kipf in testes. Typically kipf recruits the RDC complex to specific piRNA source loci throughout the genome. Data from ovary suggests that knocking out kipf leads to an enrichment of the RDC complex at satellite DNAs. B) qRT-PCR in kipf KO homozygotes (kipf/kipf) and heterozygotes (kipf/TM3) shows that complex satellites, and specifically Rsp, is upregulated in both ovaries and testes when kipf is knocked out. C) Drive strength assays show that kipf-KO individuals exhibit much lower drive strength when compared to the heterozygote in an SD-Mad background. We see a mild but significant reduction in drive in the SD-5 genotype despite not seeing a Rsp smRNA phenotype in the smRNA seq.

To determine if a loss of *kipf* (and thus overexpression of *Rsp*) is sufficient to rescue wild-type sperm from *SD-Mad,* we used a knockout generated by deleting the genomic locus (48). We examined the effects of *kipf*-KO on the drive strength of *SD-Mad* and *SD-5* in an *Iso1* background. We estimate drive strength as *k:* the proportion of the progeny of *SD* heterozygotes that contain the *SD* chromosome. A *k* value of 1 represents perfect drive and a *k* value of 0.5 represents fair Mendelian segregation. We found that knocking out *kipf* was sufficient to reduce the strength of drive in *SD-Mad* heterozygotes from ∼0.975 to ∼0.815 (*P* < 0.00001, arcsine transformation then t-test (asin-t), Figure 3C, Supplemental File 4). We observed a similar reduction in drive strength (from *k* = 0.98 to *k* = 0.88; *P* < 0.00001, asin-t, Figure S11A) when we knock down *kipf* (*kipf*-KD) compared to a white shRNA control using RNA interference (48). We also tested *kipf-*KD in a semi-sensitive *Rsp* deletion line (*Iso1*Δ*C8,* hereafter *C8*) derived from the *Iso1* chromosome but with ∼50% of the *Rsp* locus deleted using CRISPR/*Cas9* (51). In a *SD-Mad/C8* background, *kipf*-KD reduced drive from *k* = ∼0.85 to *k* = ∼0.72 (*P* < 0.00001, asin-t, Figure S11B). *Kipf-*KO had a mild but statistically significant effect on the drive of *SD-5* heterozygotes (from *k* = ∼1 to *k* = ∼0.94, *P* < 0.00001, asin-t, Figure 3C).

The reduction in drive strength in the absence of *kipf* is consistent with the hypothesis that *Rsp* transcription, transcripts, or their processed products (i.e. piRNAs) are involved in the mechanism of *SD-Mad* and are somehow important for the development of *Rsp*-bearing sperm. However, the loss of *kipf* also overexpresses other satDNAs, like *1.688* (Figure 3B), and likely dysregulates *kipf-*dependent piRNA clusters (48).

### Overexpression of Rsp piRNAs and weaker drive in SD-Mad heterozygotes

To specifically overexpress the *Rsp* repeat, we inserted 3 copies of the *Rsp* monomer into the abundantly expressed piRNA cluster, *38C1*, in an *Iso1* 2^nd^ chromosome (*Iso1^Rsp>38C1^*) (Figure 4A). This insertion results in the overexpression of *Rsp* in both *SD* backgrounds (*SD-Mad* background: Log2FC = 0.45, *P.adj* = 1.6e^-10^, DESeq2; *SD-5* Log2FC = 0.47, *P.adj* = 0.008, DESeq2) (Figure 4B-C, S12, Table S5, Supplemental File 1). We detect a mild effect on the expression of *38C1* (Figure 4B; *SD-Mad*: Log2FC = -0.34, *P.adj* = 2.3e^-11^*, SD-5*: Log2FC = -0.37, *P.adj* = 4.3e^-5^, DESeq2) but the *Rsp* insertion generally does not influence transcription at other piRNA clusters (Figure 4B).

**Figure 4:**
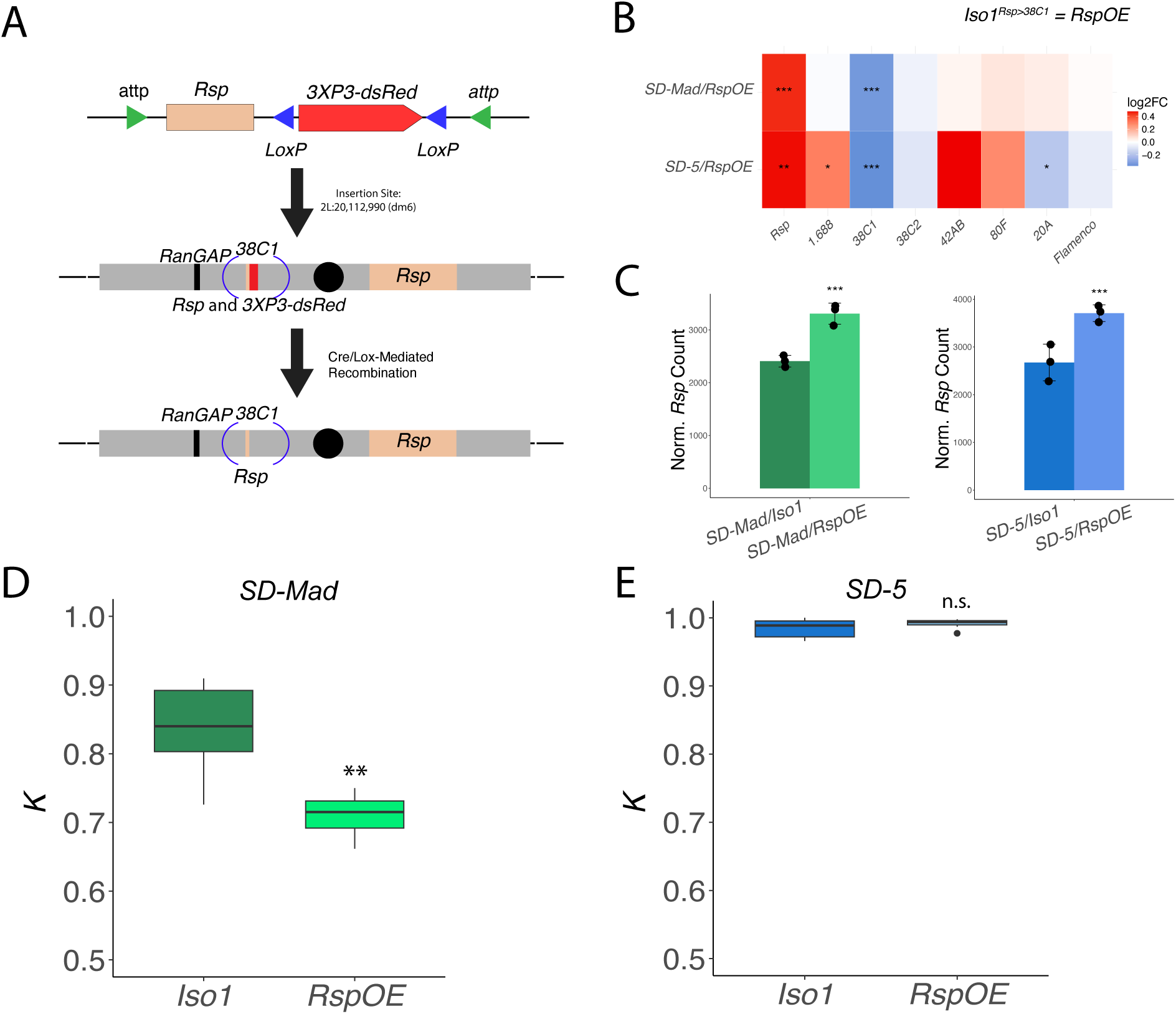
Overexpression of Rsp via 38C1 insertion is sufficient to reduce drive strength of SD-Mad heterozygotes but has no effect on SD-5 heterozygotes. A) Schematic of the overexpression construct. Four copies of Rsp were inserted roughly 1/3 of the way into the piRNA cluster 38C1 of Iso1 using CRISPR/Cas9 mediated homology directed repair. The construct included a floxed 3xP3-dsRed reporter which was removed using Cre after the insertion was validated. Attp sites were also included to facilitate additional insertions of Rsp. B) Rsp is significantly overexpressed in the RspOE strain for both SD heterozygotes relative to the same genetic background with the unmodified 2nd chromosome. 38C1 on the other hand was significantly downregulated. Other piRNA clusters were largely unaffected. C) A comparison of the DESeq2 normalized counts from SD/Iso1 and SD/RspOE smRNA. Rsp is significantly overexpressed in both conditions. D) Drive strength assay for SD-Mad in an RspOE background relative to Iso1. Drive is significantly and substantially reduced the SD-Mad/RspOE genotype. E) Drive in SD-5/RspOE was unaffected. These observations demonstrate that SD-Mad and SD-5 appear to interact with Rsp differently.

We tested the transgenic flies with the full *Rsp* overexpression construct and the 3XP3-dsRed reporter (*Iso1^Rsp-dsRed>38C1^* and did not see a change in drive strength, Figure 4A,D, S13, Supplemental File 4). We suspect that the insertion of a strong euchromatic promoter into an otherwise heterochromatic locus may affect the RNA species produced from the locus, so we removed the *3XP3-dsRed* reporter via recombination using *Cre* (Figure 4A,D, S13, Supplemental File 4).

When we tested the drive sensitivity of *Iso1^Rsp>38C1^* in an *SD-Mad* heterozygote, we found that drive strength is significantly lower relative to *Iso1* (Figure 4D, Supplemental File 4). This provides further evidence that *SD-Mad* somehow perturbs *Rsp* RNAs in the presence of *Sd-RanGAP* as part of its mechanism.

In contrast, *Rsp* overexpression had no effect on drive strength in *SD-5* heterozygotes in an otherwise identical background (Figure 4E, Supplemental File 4), consistent with the smRNA results suggesting that *Rsp* RNAs are not perturbed in *SD-5*. Taken together, these results suggest that *SD-Mad* and *SD-5* have diverged in the factors and features involved in their drive.

## Discussion

We make two major observations in our study of molecular phenotypes of *SD* testis transcriptomes. First, we show that *Rsp-*derived smRNAs are downregulated in the driving testes of *SD-Mad* heterozygotes and that overexpressing *Rsp* RNAs affects drive strength. This work therefore links RNAs complementary to the target satellite with the drive phenotype. Second, we revealed mechanistic variation in drive haplotypes—while *SD-Mad* and *SD-5* affect similar pathways, only *SD-Mad* affects *Rsp* smRNA. These results suggest that, while *SD* chromosomes share a target and main drive locus (*Sd-RanGAP*), the modifiers accumulated on each haplotype may influence the drive mechanisms, either by creating new pathways to drive or acting as tuning knobs on drive strength.

### smRNAs derived from the target satellite DNA affect drive

Our data indicate that *SD-Mad,* but not *SD-5,* disrupts the abundance of *Rsp-*derived smRNAs and that this phenotype correlates with drive. Our observations add *SD* to the growing list of meiotic drive systems that involve smRNAs (11). We observed an effect on drive strength—and the increased survival of wild-type *Rsp-*bearing sperm—even with a modest overexpression of *Rsp* RNA (∼2 fold with *kipf* KO or KD and ∼1.5-fold with *Iso1^Rsp>38C1^*). The ability of the *Iso1^Rsp>38C1^* transgene to rescue wild-type sperm may depend on transcription of the source RNAs and genetic background: we did not see an effect on drive without removing 3x*P3-DsRed* from the *38C1* insertion (Figure 4D-E, S13). The difference in drive suppression phenotypes between *Iso1^Rsp>38C1^* and *Iso1^Rsp-dsRed>38C1^* may suggest that although transcription is taking place at the *Rsp>38C1* locus in *Iso1^Rsp-dsRed>38C1^* (Figure 4B), the transcripts may not be shunted into any functional RNA pathways. Nonetheless, the *kipf* experiments (Figure 3) overexpress *Rsp* from the endogenous locus rather than a transgene in a piRNA cluster, albeit not specifically, and therefore bolsters our conclusions that *Rsp* RNAs are involved in *SD-Mad* drive. Future studies that focus on how different species of RNA might protect sperm and influence drive phenotypes may provide important clues about drive mechanisms and the functions of these RNAs.

We know relatively little about the rules that govern transcription at satellite DNAs. Previously, we found that the *Rsp* satellite is regulated by non-canonical heterochromatin-dependent transcription machinery, similar to dual-stranded piRNA clusters (34,35). There may be specific transcription factors important for licensing transcription at *Rsp,* similar to the role of *HP2* in regulating the transcription of *AAGAG* (52). Identifying potential proteins that interact with *Rsp* may therefore provide important clues about why satellites like *Rsp* are targets of drive.

Several drive systems involve smRNAs as suppressors of drive (12–14,53). *Rsp* smRNAs, on the other hand, appear to be targets of drive. The protective effect of increased *Rsp* RNAs in *SD-Mad*-mediated drive suggests that these smRNAs may play a role in chromatin regulation but these *Rsp-*derived smRNAs are not required for spermatogenesis in the absence of the *Rsp* genomic locus. Spermatogenesis normally proceeds without issue for chromosomes with *Rsp* deletions (34). Interestingly, despite being a target of *SD* drive, large *Rsp* loci are abundant in natural populations (33) and there is some evidence that there may even be a fitness advantage associated with large *Rsp* alleles, which could explain its persistence (54).

We propose a model where satDNAs require epigenetic regulation by smRNAs in a dose-dependent manner during spermatogenesis and *SD* exploits this feature to bias its transmission. The epigenetic profile of male germ cells is dynamic. Histones experience multiple post-translational modifications before ultimately being replaced by protamines (55,56). We hypothesize that the smRNAs direct H3K9 trimethylation at satDNAs. Without proper coordination, the satellite cannot be correctly repackaged with the rest of the pericentromere during meiosis or the histone to protamine transition, triggering a checkpoint on sperm chromatin quality that leads to an individualization failure. In *SD-Mad* drive, disrupting *Rsp* smRNA affects wild type locus with the large block of repeats, but *SD-Mad* chromosomes are unaffected because they lack the repeat.

Alternatively, if a lack of smRNA regulation makes *Rsp* more accessible, it may become a sink for transcription factors or other proteins. For example, the pioneering factor *GAF* binds to the simple satellite, *AAGAG,* during embryogenesis and is required for its transcriptional silencing (57). Either of these defects could be the cause for the eventual elimination of the affected spermatids. Without more knowledge on the dynamic epigenetics of the *Rsp* locus during spermatogenesis it may be impossible to tell why small RNAs are apparently necessary: at least for those spermatids which contain a large enough *Rsp* locus to be sensitive to *SD-Mad*.

It is intriguing that while *SD-Mad* involves *Rsp-*derived RNAs, we do not detect any effects of *SD-5* on *Rsp-*derived RNAs, and we see no effect on drive strength upon overexpressing *Rsp* RNA on *SD-5* drive using *Iso1^Rsp>38C1^.* This suggests that *SD-Mad* and *SD-5* drive via different specific mechanisms. It is interesting that although *SD-5* heterozygotes did not affect *Rsp* RNA, the *kipf*-KO experiments revealed a small but significant effect on drive strength. This suggests that other piRNA clusters, transposons, or satellite DNAs can affect drive. This also suggests that, while *SD-5* and *SD-Mad* have different effects on smRNAs, *SD-5* associated drive mechanisms may operate downstream of *Rsp* piRNAs.

### Mechanistic variation in drive haplotypes

We suggest that the mechanistic variation between *SD-Mad* and *SD-5* arises from molecular differences in the haplotypes. The most conspicuous differences between these and other *SD* haplotypes are in their chromosomal inversions (Figure 2A). These inversions may link different genetic variants, including enhancers of drive, to the main drive locus, creating different co-adapted gene complexes. While there is abundant genetic evidence for linked modifiers that enhance *SD* (i.e. *Enhancer of SD (E(SD))* (26)*, Modifier of SD (M(SD))* (27), and *Stabilizer of SD (St(SD))* (28) on some well-studied *SD* haplotypes, none have been molecularly identified. The suppression of recombination under chromosomal inversions and around the centromere, preventing us from mapping putative modifiers precisely. *SD-Mad* may contain a modifier that influences *Rsp* smRNA abundance that is absent in *SD-5*. Further experimentation is required to molecularly characterize the differences between these chromosomes and whether the effects on drive arise from a single locus or epistatic interactions between loci.

All *SD* chromosomes share the main drive locus, *Sd-RanGAP*, which is a duplication of an essential component of the Ran cycle (25). We suspect that there are myriad ways to induce sperm dysfunction involving the Ran cycle, and that while different *SD* haplotypes share the *Sd-RanGAP* duplication, they may recruit different modifiers. This system-level property of *SD* might affect its population dynamics. *SD* drive prompts the evolution of drive suppressors (19). *SD* chromosomes that acquire new modifiers that help escape suppression may gain a transmission advantage over *SD* haplotypes targeted by suppressors (10,33). These dynamics could explain the rapid turnover of *SD* haplotypes in populations across the globe (58–61).

### Drive-correlated transcriptional changes

Our differential expression analysis did not suggest specific modifier candidates. However, we identified potentially interesting genes among those most differentially expressed between *SD-Mad* testes with and without the main drive locus that may highlight processes involved in drive mechanisms.

First, *Tono,* a BTB zinc finger-containing transcription factor is upregulated (Log2FC^DGE^ = 1.7) in all *SD-Mad* comparisons. *Tono* plays a role in regulating transcription in muscle cells in response to mechanical pressure (62) but also shows enrichment in male germ cells (63). The putative DNA-binding capacity and ability to form nuclear condensates (62) makes this an interesting candidate gene for interacting with the *Rsp* satellite. Second, the *importin-4* ortholog, *Artemis (Arts)*, which facilitates *Ran*-mediated import of *H3* and *H4* is overexpressed in *SD-Mad* (Log2FC^DGE^ = 2.5). Interestingly, *Arts* expression is antagonistic to male fertility (64). Also of note, *Apollo,* a duplicate of *Arts* which supports male fertility (64) is downregulated (Log2FC^DGE^ = -0.6) though it is not in the top-most differentially expressed genes.

The differentially expressed genes in the transcriptomes of driving testes may be more related to the drive phenotype than the mechanism. First, we found that multiple repetitive loci are differentially expressed with little evidence that they influence drive. While this could be due to the different 2^nd^ chromosomes in the comparison, it could also be due to incidental effects from the drive mechanism on the *Rsp* locus. Second, we found that genes related to individualization and the cell cycle are broadly differentially expressed (Figure 2, S8). Third, we detect a consistent differential expression in genes involving cytoskeleton dynamics. *SD*-targeted sperm experience a delay in protamine loading and can have nuclear morphology defects (30) leading to *SD-*targeted sperm being eliminated around the individualization stage. Elongation and individualization heavily rely on actin and microtubule dynamics (65) and appears to be a common point of sperm elimination (reviewed in 45). As *SD-*mediated drive eliminates wild type sperm at the individualization stage, the enrichment of genes involved in cytoskeleton-related processes in our analyses could be related to the drive phenotype rather than its proximal cause. We suspect that the timing of the proximal cause of *SD-*mediated drive may align with early spermatogenetic processes; perhaps where cell cycle-related genes are active and appear to be broadly differentially expressed (Figure 2B, Module H). This earlier timing is consistent with temperature shift experiments that place the critical period for *SD* at or before meiosis (47).

Regardless of the precise mechanism, different *SD* chromosomes may converge on similar phenotypes if the dysfunctional sperm targeted by *SD* trigger a sperm quality checkpoint that eliminates spermatids (30,66). Many have proposed the existence of such a checkpoint. Events that take place during prophase/metaphase may be linked with late-stage sperm elimination (67,68). Lastly, a recent study identified a component of this putative checkpoint (66). The enrichment of genes related to cytoskeleton pathways found in our gene set analysis could be associated with the common elimination mechanism of this putative checkpoint. Alternatively, the *SD*-induced defect may not trigger this putative checkpoint and instead, the spermiogenic failure at individualization is simply the end point of a cascade of perturbations that begins prior to meiosis.

In this paper, we make significant progress toward understanding molecular mechanisms of *SD* in connecting *Rsp* RNA to *SD.* Future studies of *SD-*related phenotypes and effects on satellite DNA transcription and chromatin, may have general implications for understanding how spermatogenesis is vulnerable to drive and other cheaters.

Drive is widespread across species and has significant consequences for the fundamental process of gametogenesis and its evolution (reviewed in 3,69,70). Revealing drive mechanisms could help us understand fundamental aspects of spermatogenesis, with implications for our understanding of male infertility, and the development of efficacious and safe synthetic drivers to control vectors of human disease, address agricultural challenges, and species conservation.

## Methods

### Fly Stocks

We maintained all fly stocks and crosses at 25°C on cornmeal medium. *Iso1* (71), *R16* (26), *CyO/Gla*, *SD-Mad* and *SD-5* (72). *Iso1* chromosomes are typically sensitive to drive (*Rsp^s^* allele with ∼1100 copies) and *Gla* chromosomes are typically super-sensitive to drive (*Rsp^ss^* allele with >2000 copies), although the *Gla* chromosome we use here has lower drive strength in the *SD-Mad* background (Table S1)*. R16* is a complete deletion of *Rsp* and the surrounding sequences (26).

The stocks for the *kipf* knockdown experiments were provided as a generous gift from Julius Brennecke (48) and are now available from the Vienna Drosophila Stock Center: *Kipf-*sh VDRC# 314048, *w*-sh VDRC# 313772.

The *Sd-RanGAP* deletion line SD-Mad^Rev^ is a transgenic line that we made using CRISPR. We generated a gRNA-expressing construct by inserting the gRNA sequence (AGGAGGATTTGGAATAGTC) into *pCFD3* vector following the protocol in (73). This gRNA sequence matches the sequence near the breakpoint between the *Sd-RanGAP* and wild type *RanGAP* gene. We also generated a repair construct with the homology arms matching the sequence around the CRISPR cut site and a tag. To perform CRISPR on the *SD-Mad* chromosome, we generated a fly stock with the X chromosome from the Bloomington Drosophila Stock Center #51323 (*y[1] M{vas-CasS}ZH-2A w[1118]*), and the 2nd chromosome from *SD-Mad*. The injection of the *pCFD3* gRNA construct into this *Cas9;SD-Mad* fly was done by GenetiVision. The transgenic flies were screened, and the deletion of *Sd-RanGAP* was verified by PCR.

### Rsp overexpression transgenic ffy

We used the CRISPR/*Cas9* system to insert *Rsp* sequences into the *38C1* piRNA cluster to generate the *RspOE_DsRed* line. We generated a gRNA expressing construct by inserting the gRNA sequence (GATTCGATCTCAAGACCCAC) into a *pCFD3* vector following the protocol in (73). The designed gRNA targets position 20,154,890 on contig 2L_1 in the heterochromatin-enriched genome assembly (74). To generate the repair construct, we synthesized two gBlock fragments (Integrated DNA Technologies, IDT) based on the homology arms around the CRISPR cut site. We inserted the gBlock with the left homology arm into the pDsRedattp (Addgene #51019) vector with SphI and NdeI (New England biolabs, NEB), and then inserted the gBlock with the right homology arm with BglII and XhoI (NEB). We synthesized a Rsp trimer sequence by PCR using primers (forward: GGAAAATCACCCATTTTGATCGC, reverse: CCGAATTCAAGTACCAGAC) described before in (34), and inserted it into the repair construct with AvrII and SacII (NEB). All insertions were verified with Sanger sequencing. To accomplish CRISPR on the *Iso1* chromosome 2, we generated a fly stock with the X chromosome from the Bloomington Drosophila Stock Center #51323 (y[1] M{vas-Cas9}ZH-2A w[1118]), and the 2nd chromosome from *Iso1*. The injection of the pCFD3 gRNA construct and the repair construct into this *Cas9;Iso1* fly was done by GenetiVision. The transgenic flies were screened using the DsRed visible marker and we verified the DsRed insertion by PCR. To remove the DsRed marker, we crossed the RspOE_DsRed line to a Cre-expressing line y[1] w[67c23]; sna[Sco]/CyO, P{w[+mC]=Crew}DH1 (BDSC #1092) following the cross scheme described in section 3.1.3 in (75). The removal of the DsRed sequence was verified by PCR.

gBlock sequence with the left homology arm:

GCACATgcatgctagcTTAATTATTAAGTACAAAAGGAAAAATTTATCTAGTTATTAATTACTTAATGTGTAATAATATATTACTATCTTACTATCCTCCGGGTAGGGAGATCGCTGGGGTGTACCCTCTTCAGCCTTCGAAGGGAGATGGGATTAACTAGGATATTTGCCAGTCGGTTCGGGTGGACCATAAGCCTGCCGTCTGCAGAAGCAACAATCACTTCAACAACGAGAAGACAGTAATCGACAGCACAGATTTCTTGATGTAAAATCGCTTCTGGAGCAACAACAACAACAATTTCAGCAATGGCAACAGCAATTTCGCTTGTGGCTTCGACAGGAAAATCAACAGCAAAACAAACATGTCAACCAGCGCTTAAAAAAGCTTGAAATATCGTATTTGAAATCGCTAATAATATAAAACAATGAGCTGGAGCTAAACACCAGTCTCATGACAAAGTTTCGGCCTTGCAATGATCCATCTTAAGGTTCTTATATGGAGTGTCAATGGCATTTCATGCAAAGCCAGAGAAATTGAGCGCTTCGCAACTGACCTTGGTGACATGGAGTTCAGTGCCATTTACTGCCCTCCAATGAACAGATTAGAAGAAAGACATTTCACTAATCTACTCCGTGCTTGCAGGCAAAGGTACTTGGTAGATAGTGACTGGAATGCGCGACACTGGCTGTGGGGAGATAGATACAACTCACCCAGGGGGCGAGAACTAGCTAAAGCCATTTCTAGCTGTGGGGCTAATATTCTTGCAGACGTGCGATTCGATCTCAAGACCCACgcggccgcgagctcCCCAGGTCAGAAGCGGTTTTCGGGAGTAGTGCCCCAACTGGGGTAACCTTTGAGTTCTCTCAGTTGGGGGCGTAGGGCCGCCGACATGACACAAGGGGTTGTGACCG GGGTGGACACGTACGCGGGTGCTTACGACCGTCAGTCGCGCGAGCGCGAcctaggTCCccgcggAGGCATTcatatgGAATTCC

gBlock sequence for the right homology arm:

GGAagatctggcgcgccTCGCGCTCGCGCGACTGACGGTCGTAAGCACCCGCGTACGTGTCCACCCCGGTCACAACCCCTTGTGTCATGTCGGCGGCCCTACGCCCCCAACTGAGAGAACTCAAAGGTTACCCCAGTTGGGGCACTACTCCCGAAAACCGCTTCTGACCTGGGaggcctgtcgacggtaccATGACTTCTTTGGAACCGGGCGGCAAATCAGCTAAAAAACTTGCTAAGACTTAGAAACGAATTCTTTGAGCAAAAGTTGACGTCGCTGGACTAGACTAGTGGGCATAAGTATTTGGACAAGAAATATTTTCAATATATTTGTTTGTAACTCATTTTTTACTTAAAGCAAAATTGGTTAATTATTTTGTTGATTTAACAAAAATTACTTTCTACAAATTAGCAGTGGCATAAGTACTTGGACAAATTATTTAAATGTTATAAATTGAAAAAAAAAGTCAACTACAAAATTATGACAAATTAATAAAAAATATACACTCCCTCAGCGTCTATCACCTTTTGCAGACGTTTTGGTACAGACTGTACCAAGCTGTGGATATAATAATCTATAATCTATAATCTATTGAAATATCCTTCCATAAGGTTTGAATTTCCAAAATCGTTTATTCGCGACTTTTAGATAAGTTTGCACTCCATTTTCTTTTTAAATATGACCAGAGGTTTTCTATTATGTTGAGGTCTGGACTTTGAGGTGGCCAATCCAATGTGTTTACGCCAACATCTTTCAGAAACTTCGTAATCAGTTTGGCCTTATGACAGGGAGCATTATCCTGCTGGAGTATGAAGGACTCTCCAATCAATTTATCACCAGAGGGGAATGCATGATTATTAAGGACATCTAAATATTTTCCTTGATTCATGGTTCCATCAACAGGAACCAAATCTCCCAAGCCAGTGAAAGGATATTTTGCTGTTCACAAAGCTTCTTATAAGCAAATATTGGTTTTTAGACgctcgaggggcgccAAA

*Rsp* PCR product sequence:

CCGAATTCAAGTACCAGACAAACAGAAGATACCTTCTACAGATTATTTAAACCTGGTACACAAAAACAATAAATTGACTAAGTTATGTCATTTTAAGCGGTCAAAATTGGTGATTTTTCGATTTCAAGTACCAGGCGAACAGAAGACACCTTCTAGAGATTCTGTTCAACTGGTAAGCAAAAACAGTAAATTGCCTAAGTTTTACATTTTAAGCGGTCAAAATGGGTGATTTTCCGATTTCAAGTACCAGACAAACAGAAGATACCTTCTACAGATTATTTAAACCTGGTACACAAAAACAATAAATTTACTAAGTTATGTCATTTTAAGCGATCAAAATGG GTGATTTTCC

### Testis RNA Extraction for (q)RT-PCR

10-15 pairs of testes per replicate from 3-5 day old flies were dissected in cold PBS and immediately placed in 300ul of 1X RNA Protect from the Monarch Total RNA Miniprep kit (New England Biolabs Cat # T2010S). The tissues were stored in a -80C freezer until all replicates were dissected. When RNA was to be isolated, we added proteinase K and buffer to the sample and incubated the testes at 55C for 20-30 minutes. Genomic DNA was removed using a gDNA removal column and one volume of 95% EtOH was added to the eluent (the fraction containing the RNA). The RNA was run through an RNA capture column and was subsequently treated with on-column *DNAse I* for 45-60 mins at RT (extended DNase treatment allows more time to degrade repetitive DNA). The RNA was then primed with RNA Prime buffer and washed twice. After the second wash, the tubes were spun for an additional 2 minutes to dry the membrane. RNA was then eluted in 40-50 ul of RNase free water.

### cDNA Generation for RT-PCR

200 ng of total RNA mixed with oligo(dT), satDNA-specific primer, and nucleotides was brought up to 13 ul with water and incubated at 65C for 5 minutes and allowed to cool on ice for at least 1 min. After cooling the RNA, we added 4 ul 5X reaction buffer, 1 ul DTT, 1 ul RNaseOUT™ Recombinant Ribonuclease Inhibitor (ThermoFisher Cat # 10777019), and 1 ul SuperScript™ III reverse transcriptase (ThermoFisher Cat # 18080093) (water was added for no RT controls) up to 20 ul total reaction volume. The reaction was then incubated at 25C for 5 min, 55C for 1 hr, and 75C for 15 min. If the sample was going to be used for RT-PCR the cDNA was frozen at -80C until use. If the cDNA was to be used for qRT-PCR it was diluted 3X by adding 40 ul RNase-free water and then stored at -80C until use.

### RT-PCR

1 ul of cDNA was added to 5 ul Q5® High-Fidelity DNA Polymerase rxn buffer, 2.5 ul of dNTPs, 2.5 ul of 6 uM primers, 13.5 ul of water, and 0.5 ul of Q5® High-Fidelity DNA Polymerase (New England Biolabs Cat # M0491S) for a total reaction volume of 25 ul. The samples were then cycled between 98C, 60C, and 72C 38 times and the amplicons were run on a DNA gel. Primers for kip: FWD: ACCAGCGAAACGGGATTATTA REV: CAAGGCGTAGACACTACTGAAC and RP49: FWD: GCTTCTGGTTTCCGGCAAGGTATGT REV: ACGTTTACAAATGTGTATTCCGACCACGTT.

### qRT-PCR

4 ul of cDNA was added to 7.5 ul of PerfeCTa SYBR Green FastMix (Quantabio Cat # 95072-012), 1.5 ul 6 mM primers (Rsp. For: GGAAAATCACCCATTTTGATCGC Rev: CCGAATTCAAGTACCAGAC, 359bp. For: TACGATCTCAGCGAGGTATGA Rev: TTCCAAATTTCGGCCATCAAAT), and 2 ul water for a reaction volume of 15 ul. The reactions were then run on a BioRad C1000 thermal cycler (BioRad Cat # 1851196) equipped with a BioRad CFX96 attachment (BioRad Cat # 1845097). The program consists of a 98C melting temp, 60C annealing temp, and a 10 second extension at 72C.

qRT-PCR data were analyzed using the ΔΔCt method (76) and represented using fold change (FC) relative to a housekeeping gene (*rbp4.* For: CAATCCCAACTACAATCCCTATCT Rev: CTTGGGTGCCATCGGAATTA or *rps3*. For: AGTTGTACGCCGAGAAGGTG Rev: TGTAGCGGAGCACACCATAG).

### RNA FISH

To image *Rsp* and *1.C88* we used a custom Stellaris probe set for the former and a previously described probe labeled with Cy5, (5’-Cy5TTTTCCAAATTTCGGTCATCAAATAATCAT-3’) (77) for the latter.

Testes were dissected in cold PBS and fixed using 4% FA in PBST (0.1% Tween or Triton, either produced good results) for 30 minutes on a nutator at RT. After the fixative was removed, the tissues were washed with 1 ml PBS twice, 5 minutes each on a nutator. The tissues were prehybridized for 5 minutes using 2X SSC with 10% formamide. The probe was diluted in Stellaris hybridization buffer mix (final concentration of probes: 0.1 uM for oligo probes and 0.125 uM for stellaris probes) to the samples and placed them at 37C on a nutator overnight.

After overnight incubation we washed the tissues with 1 ml of 2X SSC with 10% formamide for 30 min twice at 37C. The tissues were then mounted using SlowFade Diamond Antifade mountant (ThermoFisher, cat # S36963). The DAPI was allowed to soak into the tissue overnight at 4C before imaging within 1 week.

### K value assays

To assay the strength of segregation distortion k values were estimated from heterozygous *SD*/*Rsp^s^* males (where *Rsp^s^* are chromosomes with sensitive *Rsp* alleles) as described previously (78). At least 10 replicate crosses of single 3 to 5-day old males with two 3 to 5-day old Iso-1 female virgins were prepared. Each cross was transferred to a fresh food vial after 3 days, and the parents discarded after 4 more days. We counted the progeny from each vial through day 18 after the parents were first introduced to the vial. Only replicates with more than 50 total progeny were included in our analysis. The transmission rate of the *SD* allele depends on not only its distortion strength, but also its viability relative to the other allele. Thus, to correct for viability effects, the transmission rate of the SD chromosome was measured in *SD*/*Rsp*^s^ females. The *SD*/*Rsp*^s^ females have a Mendelian segregation ratio with k = 0.5, and any distortion from this equal segregation can be attributed to the difference in the relative viability of the two alleles. 3 replicate crosses of five 3 to 5-day old females with three 3 to 5-day old *Iso1* males were prepared, and then transferred to a fresh food vial after 4 days and kept for 4 more days. The progeny from each vial were then counted through day 18 after the parents were first introduced to the vial.

The relative viability factor w = n*^SD^* /n*^Iso1^* for each *SD*/*Rsp^s^* genotype, where n*^SD^* and n*^Iso1^* are the numbers of *SD* and *Iso1* bearing progeny produced by the *SD/Rsp*^s^ female, respectively. This viability factor was then used to calculate a corrected strength of segregation distortion k* = N*^SD^* / (*w* * N*^Gla^* + N*^SD^*), where N*^SD^* and N*^Iso1^* are the numbers of *SD* and *Iso1* bearing progeny produced by the *SD/Rsp*^s^ male, respectively. The viability corrected *k* value

(*k*\*) was reported where we were able to estimate viability (Table S1; Fig 4, S11, S12), otherwise the reported value is *k* (Fig 3).

### Imaging

A Leica SP5 laser scanning confocal microscope was used to acquire images of our slides.

### Small RNA Seq

Six- to eight-day-old testes were dissected in RNase free PBS buffer. Total RNA was extracted using mirVanaTM miRNA Isolation Kit (Ambion) with procedures for isolating RNA fractions enriched for small RNAs (<200nt), then treated with RNase free DNase I (Promega) at 37°C for 1 hour. Library preparation and sequencing were performed by Genomics Research Center at University of Rochester. Briefly, 2S rRNA was depleted (79), small RNA library was prepared with TruSeq Small RNA Library Prep Kit (Illumina) and sequenced by Illumina platform HiSeq2500 Single-end 50 bp.

### Total RNA-seq

Six- to eight-day-old testes were dissected in RNase free PBS buffer. Total RNA was extracted using mirVanaTM miRNA Isolation Kit (Ambion) with procedures for isolating RNA fractions enriched for long RNAs (>200nt), then treated with RNase free DNase I (Promega) at 37°C for 1 hour. Library preparation and sequencing were performed by Genomics Research Center at University of Rochester. Briefly, rRNA was removed and total RNA library was prepared with TruSeq Stranded Total RNA Library Prep Human/Mouse/Rat (Illumina) and sequenced by Illumina platform HiSeq2500 Paired end 125 bp. Care was taken when extracting all RNA samples that all chromosomes outside of the 2^nd^ were controlled for to control for copy number of other repetitive elements.

### Total RNA Seq Analysis

Fastq files were trimmed using trimgalore with a minimum read length of 20bp. FastQC was used to assess the quality of the remaining reads. Total RNA reads were mapped to a heterochromatin enriched genome using both Bowtie2 (for repeats) (80) and STAR (for genes) (81) and expression was calculated using HTSeq (for genes) (82) or a custom proportional counting script (for repeats; https://github.com/LarracuenteLab/SD-RNAseq). To calculate differentially expressed genes we used DESeq2 (83). We additionally used PANGEA (43) to find enriched gene sets.

Weighted Gene Correlation Network Analyses (WGCNA) (46) was carried out using the WGCNA package in R using the 50^th^ percentile of variable genes and a minimum R^2 value of 0.8. We used the spearman method for calculating correlations then found the difference between the non-driving genotype (*R16/Gla*) and the driving genotypes (*SD-Mad/Gla, SD-5/Gla*). We calculated significance using a t-test with Tukey’s multiple hypothesis correction. We then performed PANGEA gene set analysis on the modules correlating to the drive and SD haplotype. Gene sets were labeled significant if their Benjamin & Hochberg false discovery rate <0.05.

### Small RNA Seq Analysis

SmRNA seq fastq files were trimmed for quality and length with a minimum length of 18 and a maximum length of 32 using cutadapt. Remaining reads were then mapped to a library of rRNAs and tRNAs and unmapped reads were saved to a separate file. The unmapped reads were then mapped to a heterochromatin enriched genome using Bowtie (84). Counts for miRNAs and complex satellite repeats were calculated using a custom python script that assigned reads to features according to the proportion of the read mapping to that feature (https://github.com/LarracuenteLab/SD-RNAseq). Differential expression was then calculated using DESeq2 (83).

## Acknowledgements

This work was supported by an NIH R35 GM119515 to AML, an NIH F31 HD121176 to LTE, and a University of Rochester Messersmith Fellowship to XW. We thank members of the Larracuente lab for feedback on the project and the manuscript. Special thanks go to Artemis Shung, Gefei Yu, and Natasha Kruelski for discussions related to experiments in the manuscript. We thank Julius Brennecke for providing the fly lines used in our *Kipferl* experiments. We thank Nathaniel and Helen Wisch for their support. Finally, we are grateful for access to the Center for Integrated Research Computing and the Genomics Research Center at the University of Rochester, for access to computing resources and sequencing services, respectively.

## Data availability statement

All genomic data are being deposited in NCBI’s short read archive under the accession PRJNA1375835. All code and data underlying analyses in the manuscript will be available in Dryad (https://doi.org/10.5061/dryad.v9s4mw792) and code are available on GitHub (https://github.com/LarracuenteLab/SD-RNAseq).

## Supplemental Tables

**Table S1.** Estimated drive strength for *SD* and "wild type" chromosomes used in the RNAseq analysis.

| Genotype | *SD*<br>chromosome | *Rsp<sup>s</sup>*<br>chromosome | male transmission |  |  | female<br>transmission |  |
| --- | --- | --- | --- | --- | --- | --- | --- |
|  |  |  | *k<sub>m</sub>*<br>±SEM | *k<sub>m</sub><sup>\*</sup>*<br>±SEM | *n* | *k<sub>f</sub>*±SEM | *n* |
| *SD-5/Gla* | *SD-5* | *Gla* | 1<br>±0.000 | 1<br>±0.000 | 1515 | 0.518<br>±0.017 | 685 |
| *SD-5/Iso-1* | *SD-5* | *Iso-1* | 1<br>±0.000 | 1<br>±0.000 | 1057 | 0.533<br>±0.025 | 973 |
| *SD-Mad/Gla* | *SD-Mad* | *Gla* | 0.875<br>±0.024 | 0.848<br>±0.029 | 1026 | 0.560<br>±0.020 | 473 |
| *SD-Mad/Iso-1* | *SD-Mad* | *Iso-1* | 0.984<br>±0.005 | 0.981<br>±0.006 | 1633 | 0.549<br>±0.012 | 1548 |
| *SD-Mad<sup>Rev</sup>/Iso-1* | *SD-Mad<sup>Rev</sup>* | *Iso-1* | 0.582<br>±0.012 | 0.541<br>±0.012 | 1346 | 0.541<br>±0.011 | 884 |

**Table S2.** Differential expression data for piRNA clusters in the small RNA comparisons for *SD-Mad*, *SD-5*, or *SD-Mad^Rev^* with an *Iso1* or *Gla* 2nd chromosome.

|  | SD-Mad |  |  |  | SD-MadRev |  | SD-5 |  |  |  |
| --- | --- | --- | --- | --- | --- | --- | --- | --- | --- | --- |
|  | Iso1 |  | Gla |  | Iso1 |  | Iso1 |  | Gla |  |
|  | Log2FC | p.adj | Log2FC | p.adj | Log2FC | p.adj | Log2FC | p.adj | Log2FC | p.adj |
| 1.688 | -1.10 | 4.78E-14 | -0.28 | 0.001 | -0.63 | 1.01E-17 | -0.71 | 0.024 | 0.05 | 0.489 |
| *Rsp* | -0.95 | 5.09E-09 | -0.68 | 2.57E-14 | -0.21 | 0.214 | -0.07 | 0.796 | 0.33 | 2.32E-11 |
| 20A | -0.05 | 0.558 | 0.39 | 7.29E-08 | 0.02 | 0.612 | -0.14 | 0.491 | 0.23 | 3.83E-06 |
| *flamenco* | -0.18 | 0.023 | -0.14 | 0.066 | -0.14 | 0.048 | -0.45 | 9.82E-08 | -0.30 | 4.53E-13 |
| 42AB | -0.08 | 0.425 | -0.45 | 1.20E-11 | -0.15 | 0.020 | -0.16 | 0.187 | 0.07 | 0.343 |
| 38C1 | 0.22 | 0.020 | 0.16 | 0.060 | -0.05 | 0.499 | -0.03 | 0.773 | -0.09 | 0.085 |
| 38C2 | 0.60 | 1.02E-12 | 0.44 | 4.69E-09 | 0.49 | 1.49E-08 | 0.64 | 0.000 | 0.34 | 1.24E-13 |
| 80F | -0.22 | 0.100 | -0.37 | 0.003 | -0.27 | 1.70E-05 | 0.08 | 0.770 | 0.01 | 0.888 |

**Table S3.** Differential expression data for piRNA clusters in the total RNA comparisons for *SD-Mad* and *SD-5* with *Iso1* or *Gla* 2nd chromosome.

|  | SD-Mad |  |  |  | SD-5 |  |  |  |
| --- | --- | --- | --- | --- | --- | --- | --- | --- |
|  | Iso1 |  | Gla |  | Iso1 |  | Gla |  |
|  | Log2FC | p.adj | Log2FC | p.adj | Log2FC | p.adj | Log2FC | p.adj |
| 1.688 | -0.19 | 1.60E-07 | -0.19 | 1.91E-05 | 0.32 | 2.05E-03 | -0.08 | 0.149 |
| *Rsp* | -0.04 | 0.922 | 0.85 | 4.10E-03 | -0.89 | 0.146 | 0.09 | 0.821 |
| 20A | 0.35 | 0.327 | -0.09 | 0.658 | 0.90 | 0.117 | -0.22 | 0.557 |
| *flamenco* | 0.37 | 0.031 | 1.40E-03 | 0.994 | -0.28 | 0.429 | -0.37 | 0.26 |
| 42AB | -0.20 | 0.280 | 0.47 | 0.170 | 1.85 | 0.000 | -0.01 | 0.980 |
| 38C1 | 0.26 | 0.196 | 0.08 | 0.829 | 0.11 | 0.371 | -0.50 | 0.041 |
| 38C2 | 0.33 | 0.001 | -0.01 | 0.99 | 0.30 | 0.241 | -0.77 | 0.037 |
| 80F | 0.31 | 0.196 | 0.98 | 1.37E-13 | 0.35 | 0.340 | 0.42 | 5.42E-07 |

**Table S4.** Difference in Spearman Correlation coefficients for individual WGCNA modules between *SD-5/Gla* or *SD-Mad/Gla* versus *R1C/Gla*.

| Module | Comparison | Difference | P.Adj |
| --- | --- | --- | --- |
| Module A | *SD5-R16* | 0.3798 | 0.001 |
| Module A | *SDMad-R16* | 0.8058 | 1.18E-05 |
| Module A | *SDMad-SD5* | 0.4260 | 0.0005 |
| Module B | *SD5-R16* | -0.7354 | 3.79E-06 |
| Module B | *SDMad-R16* | -0.6629 | 7.20E-06 |
| Module B | *SDMad-SD5* | 0.0725 | 0.245 |
| Module C | *SD5-R16* | -0.6931 | 2.61E-05 |
| Module C | *SDMad-R16* | 0.0109 | 0.976 |
| Module C | *SDMad-SD5* | 0.7039 | 2.38E-05 |
| Module D | *SD5-R16* | 0.0756 | 0.970 |
| Module D | *SDMad-R16* | -0.1565 | 0.878 |
| Module D | *SDMad-SD5* | -0.2321 | 0.757 |
| Module E | *SD5-R16* | 0.3470 | 0.496 |
| Module E | *SDMad-R16* | 0.3559 | 0.480 |
| Module E | *SDMad-SD5* | 0.0089 | 0.999 |
| Module F | *SD5-R16* | -0.2334 | 0.036 |
| Module F | *SDMad-R16* | 0.5445 | 0.0006 |
| Module F | *SDMad-SD5* | 0.7779 | 7.69E-05 |
| Module G | *SD5-R16* | 0.0241 | 0.921 |
| Module G | *SDMad-R16* | -0.6825 | 8.19E-05 |
| Module G | *SDMad-SD5* | -0.7065 | 6.71E-05 |
| Module H | *SD5-R16* | 0.8122 | 7.32E-07 |
| Module H | *SDMad-R16* | 0.4346 | 2.12E-05 |
| Module H | *SDMad-SD5* | -0.3776 | 4.86E-05 |
| Module I | *SD5-R16* | -0.0790 | 0.969 |
| Module I | *SDMad-R16* | -0.0192 | 0.998 |
| Module I | *SDMad-SD5* | 0.0598 | 0.982 |

**Table S5.**
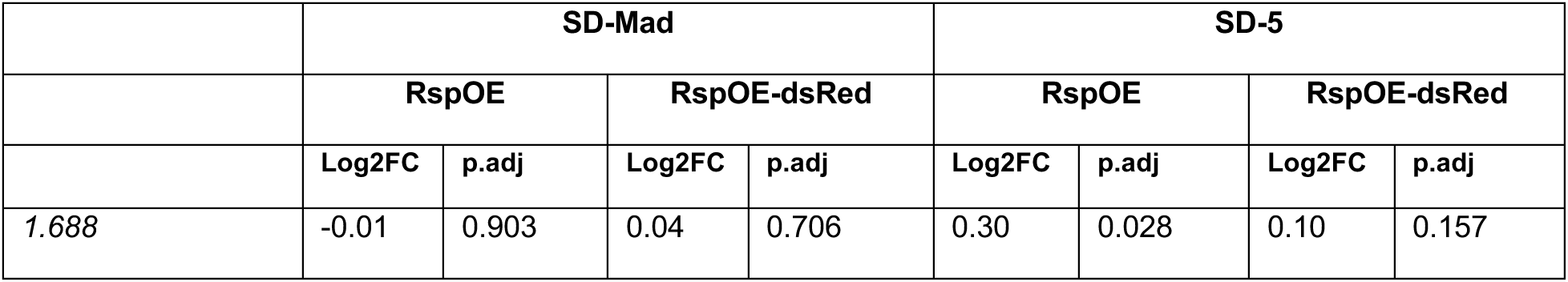
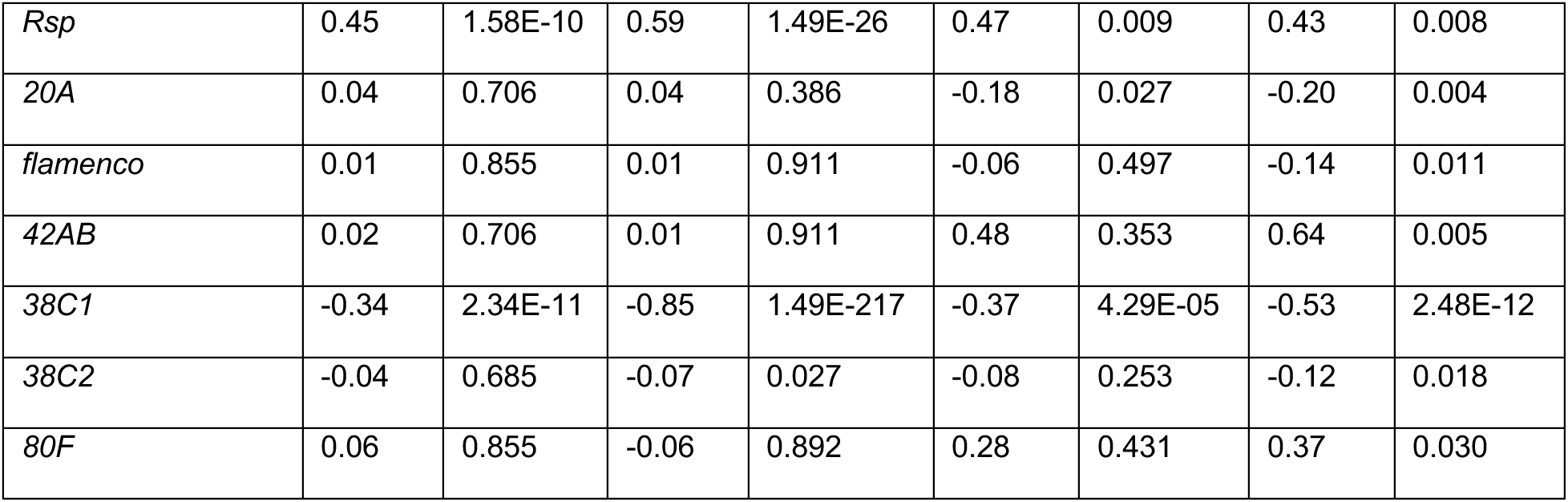
Differential expression data for piRNA clusters in the small RNA comparisons for *SD-Mad* and *SD-5* paired with either *RspOE* or *RspOE-dsRed* compared to *SD-Mad/Iso1* or *SD-5/Iso1*.

## Supplemental Figures

**Supplemental Figure 1:**
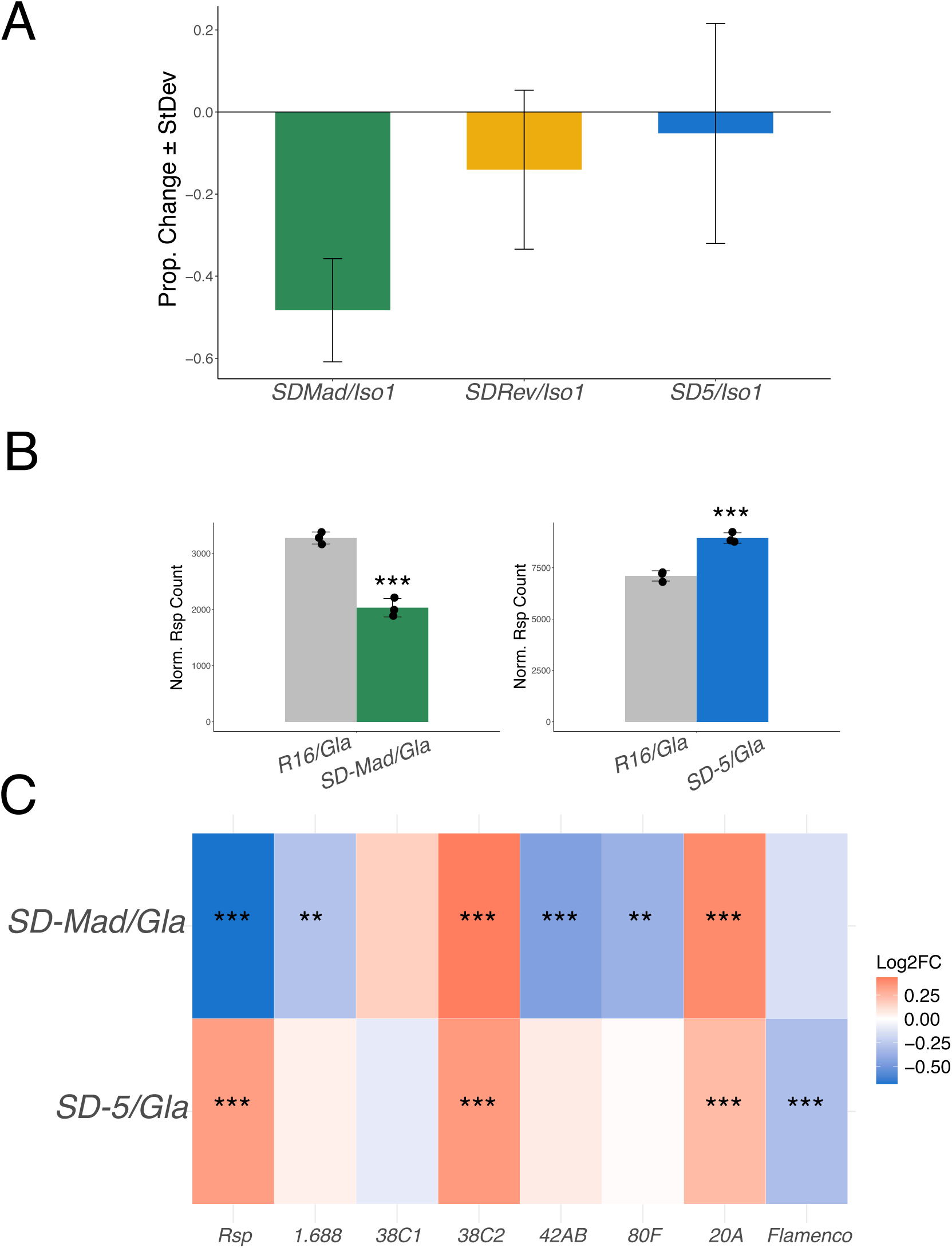
Small RNA seq analysis in *SD/Gla* backgrounds shows that small RNAs corresponding to *Rsp* are significantly less abundant in *SD-Mad* but not *SD-5*. A) A bar chart showing the change in *Rsp* small RNAs in proportion to *R16/Iso1*. *SD-Mad/Iso1* has ∼50% fewer *Rsp* smRNAs than expected. *SD-MadRev/Iso1* and *SD-5/Iso1* have largely unchanged levels of *Rsp* relative to *R16/Iso1*. B) Bargraphs showing the DESeq2 normalized counts of reads corresponding to *Rsp*. *SD-Mad/Gla* has significantly less Rsp than the control. *SD-5* has a larger count than expected. The apparent overexpression of *Rsp* in the *SD-5/Gla* condition is likely due to an epistatic interaction ectopically overexpressing *Rsp* and not drive because in all *SD/Gla* backgrounds *Rsp* is slightly higher than expected relative to *R16/Gla*. C) A heatmap showing the magnitude and significance of differential expression for *Rsp*, *1.C88*, and major piRNA clusters. *38C2* is linked to *SD* and its upregulation may be due to an epistatic interaction in the background as there is no evidence that *38C2* is involved. The somatic cell specific piRNA cluster on the X-chromosome, *20A*, is also upregulated, but it is unclear if or how this is related to drive.

**Supplemental Figure 2:**
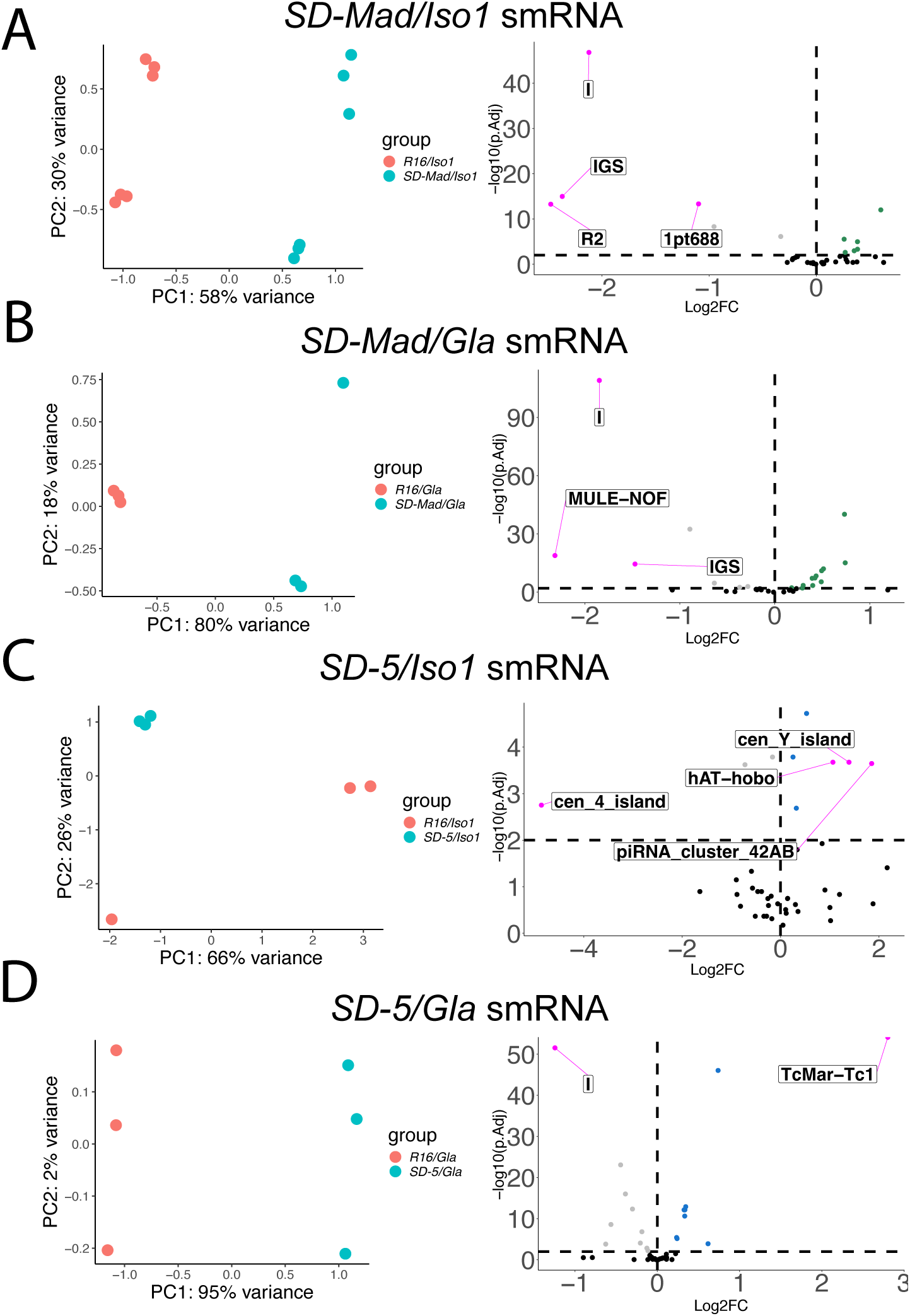
The small RNA seq samples segregate by genotype in the DESeq2 repeat analysis. A-D) PCA plots that show the two highest principle components contributing to variance between the small RNA samples in the repeat analysis. The samples largely segregate by genotype suggesting that the bulk of the variance between samples comes from the 2nd chromosomes used in the study. It is important to note that the samples in the *SD-Mad/Iso1* backgrounds come from two different batches, but that the largest contribution to the variance between replicates is the second chromosome used in the experiment. Also, one replicate from the *SD-5/Iso1* comparison didn’t group well with the other two replicates which reduces our power in this comparison. The top 5 most differentially expressed repeat elements are labeled with their common name.

**Supplemental Figure 3:**
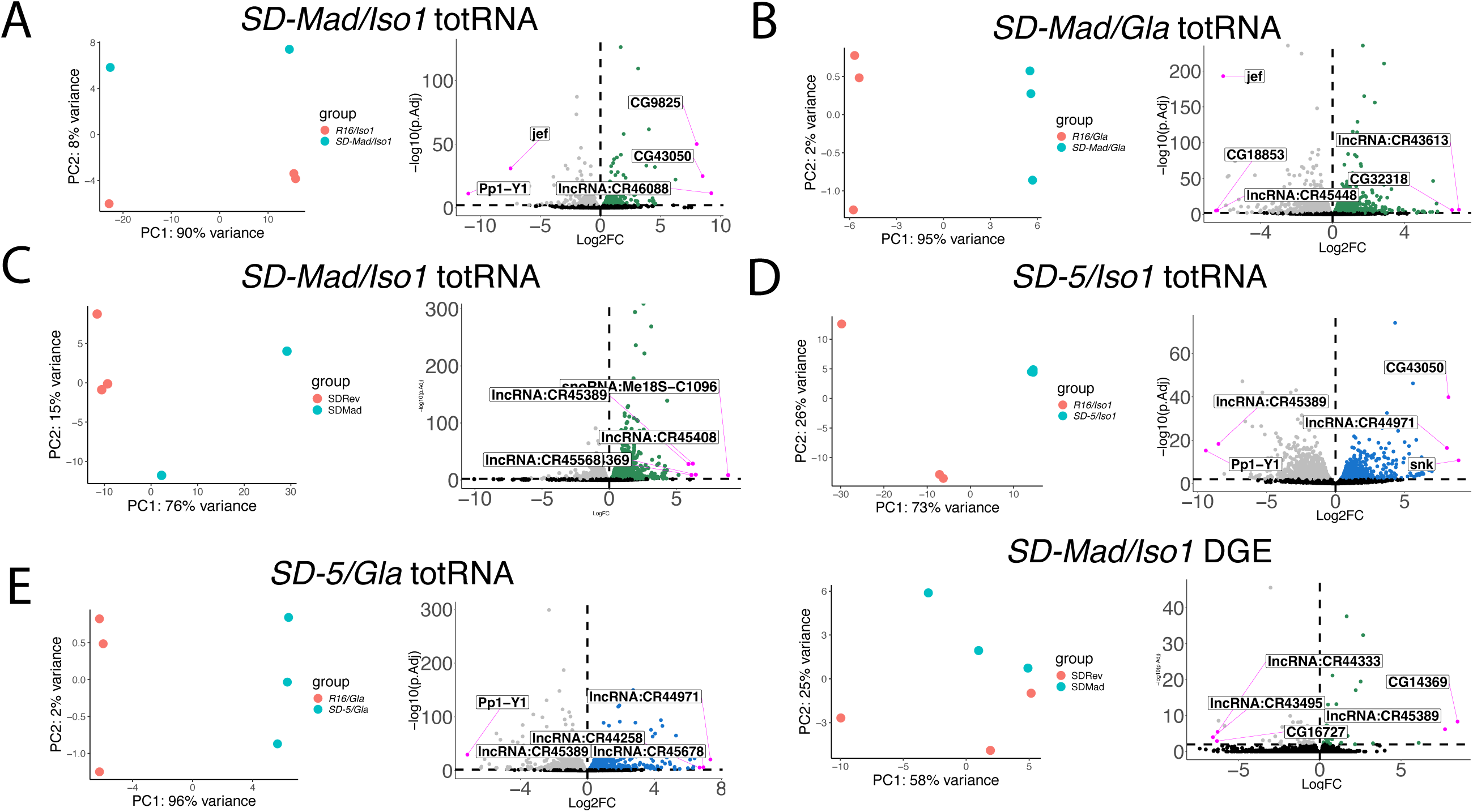
The total RNA seq samples segregate by genotype in the DESeq2 gene analysis. A-E) PCA plots that show the two highest principle components contributing to variance between the total RNA samples for the gene analysis. The samples largely segregate by genotype suggesting that the bulk of the variance between samples comes from the 2nd chromosomes used in the study. It is important to note that the *SD-Mad/Iso1* replicates have a lot of variability which limits the statistical power of comparisons with *SD-Mad/Iso1*. The volcano plots show the distribution of differentially expressed genes for each comparison with colored dots (green for *SD-Mad* comparisons, blue for *SD-5* comparisons) being upregulated relative to the copy number control (*R16/Iso1* or *Gla*) with a filter of P<=0.01 while the gray dots are downregulated. The 5 genes with the greatest Log2 fold change are labeled with the genes common name.

**Supplemental Figure 4:**
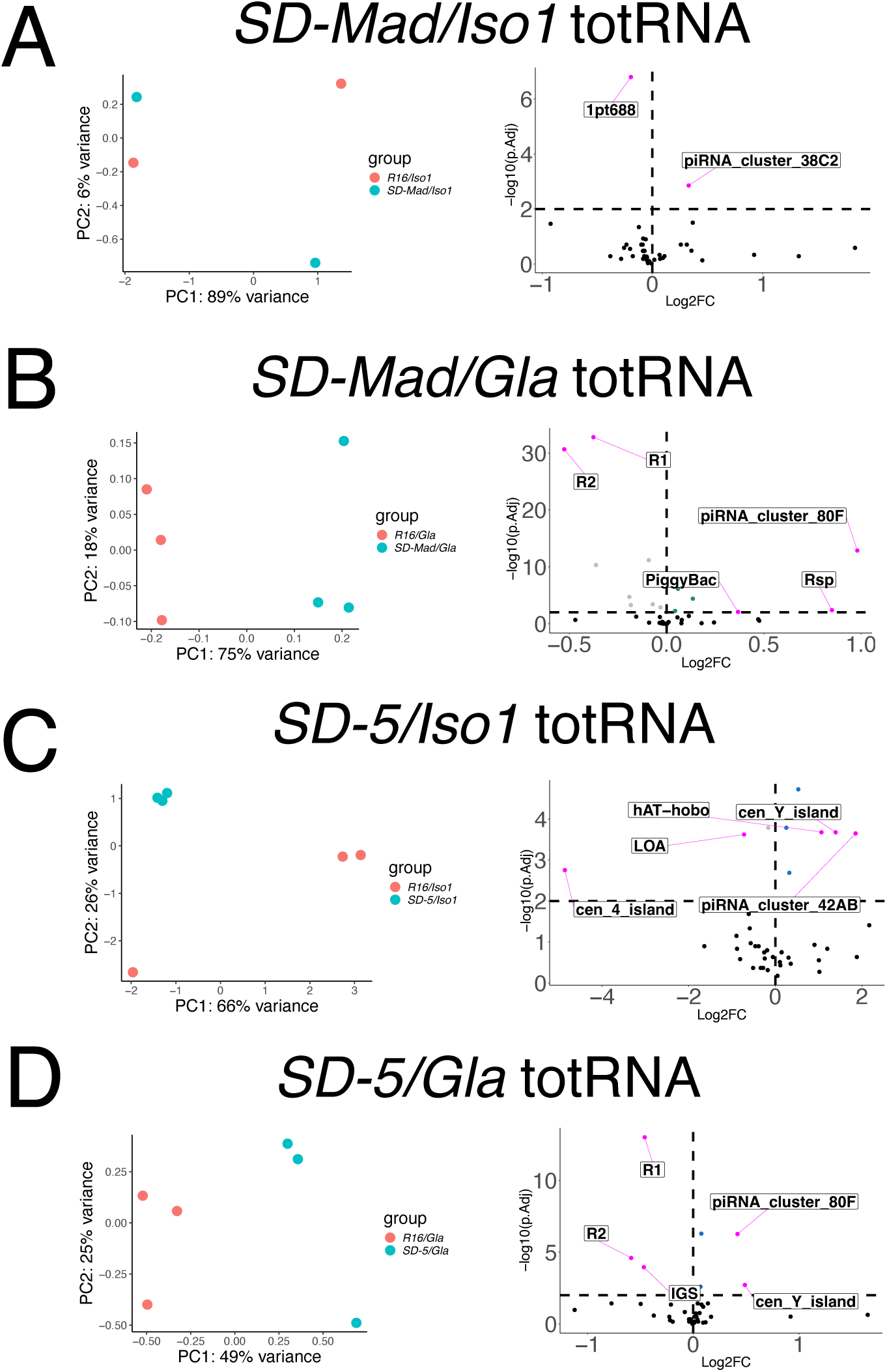
The total RNA seq samples segregate by genotype in the DESeq2 Repeat analysis. A-D) PCA plots that show the two highest principal components contributing to variance between the total RNA samples for the repeat analysis. Both *SD/Gla* backgrounds segregate well by genotype. The *SD/Iso1* comparisons do not cluster well by genotype. This limits the statistical power for the *SD/Iso1* comparisons. The volcano plots show the distribution of differentially expressed repeats as defined by DESeq2 for each comparison. Colored dots are upregulated features with a filter of P=0.01 while the gray dots are downregulated relative to the copy number control (*R1C/Iso1* or *Gla*). The 5 features with the greatest Log2 fold change are labeled with the repeats common name.

**Supplemental Figure 5:**
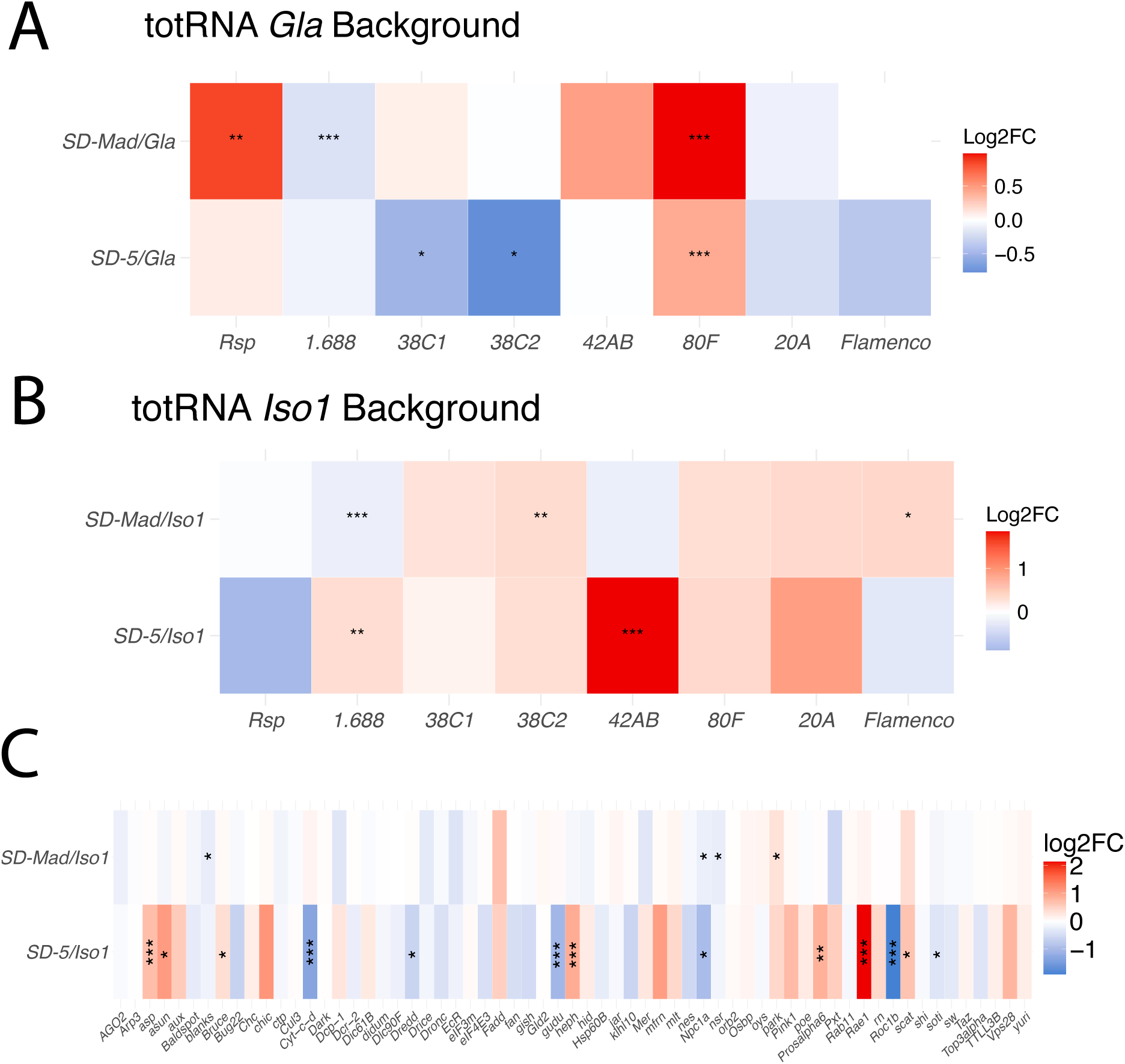
There are no patterns of differential expression in repetitive elements in the total RNA but a broad dysregulation of genes known to cause an individualization defect. A-B) Heatplots showing the significance and magnitude of differential expression for two satellite DNAs (*Rsp* and *1.C88*) and other major piRNA clusters. Rsp precursors appear to be upregulated in the *SD-Mad/Gla* background. This could be due to an epistatic interaction between the *SD* chromosomes and the other chromosomes in the *Gla* crosses. Additionally, the Chr 3 linked piRNA cluster *80F* is upregulated in both *SD/Gla* heterozygotes. C) A heatmap of differentially expressed genes in a set of genes known to cause individualization defects when dysregulated. Fewer genes are significant in this comparison than in the *SD/Gla* background. This is likely due to reduced power in the Iso1 comparisons.

**Supplemental Figure 6:**
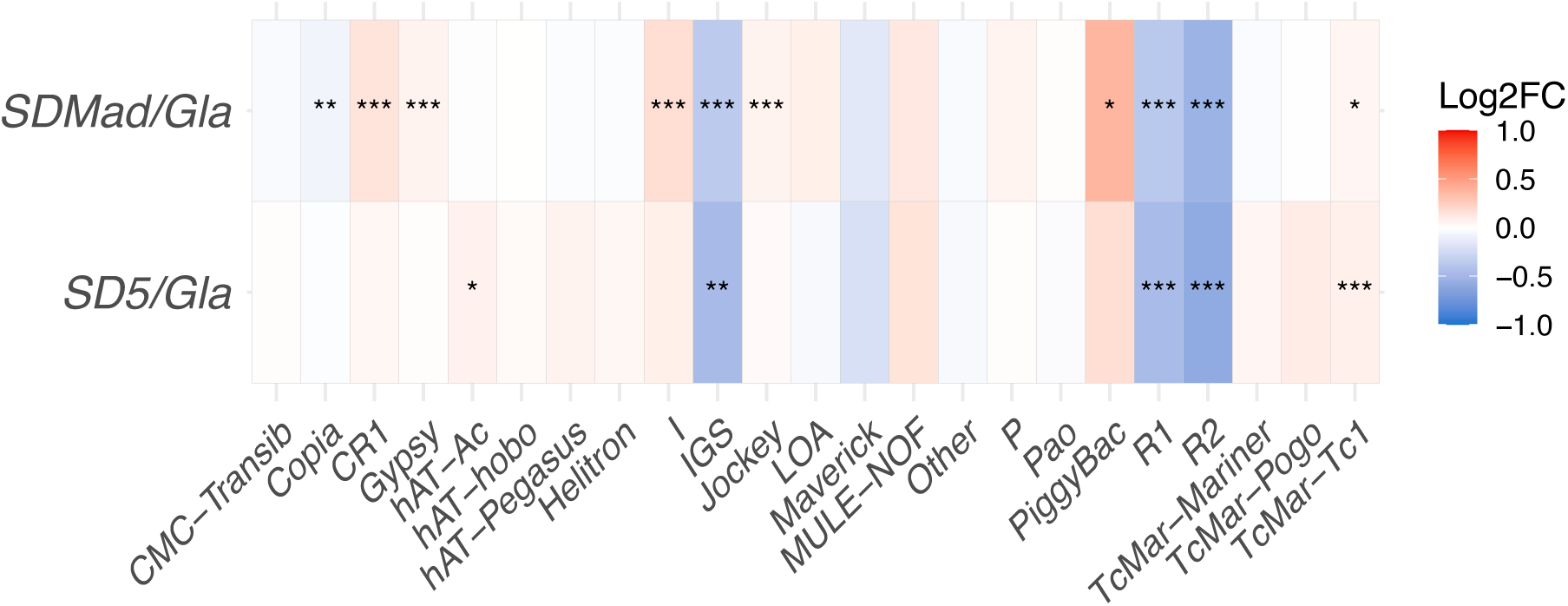
Few TE families are consistently and significantly differentially expressed in *SD* backgrounds. The heatplot shows the magnitude and significance of the change in the expression of TE families. Notably *R1*, *R2*, and ribosomal intergenic spacers are significantly less abundant than expected relative to the control. *TcMar-Tc1* appears to be upregulated.

**Supplemental Figure 7:**
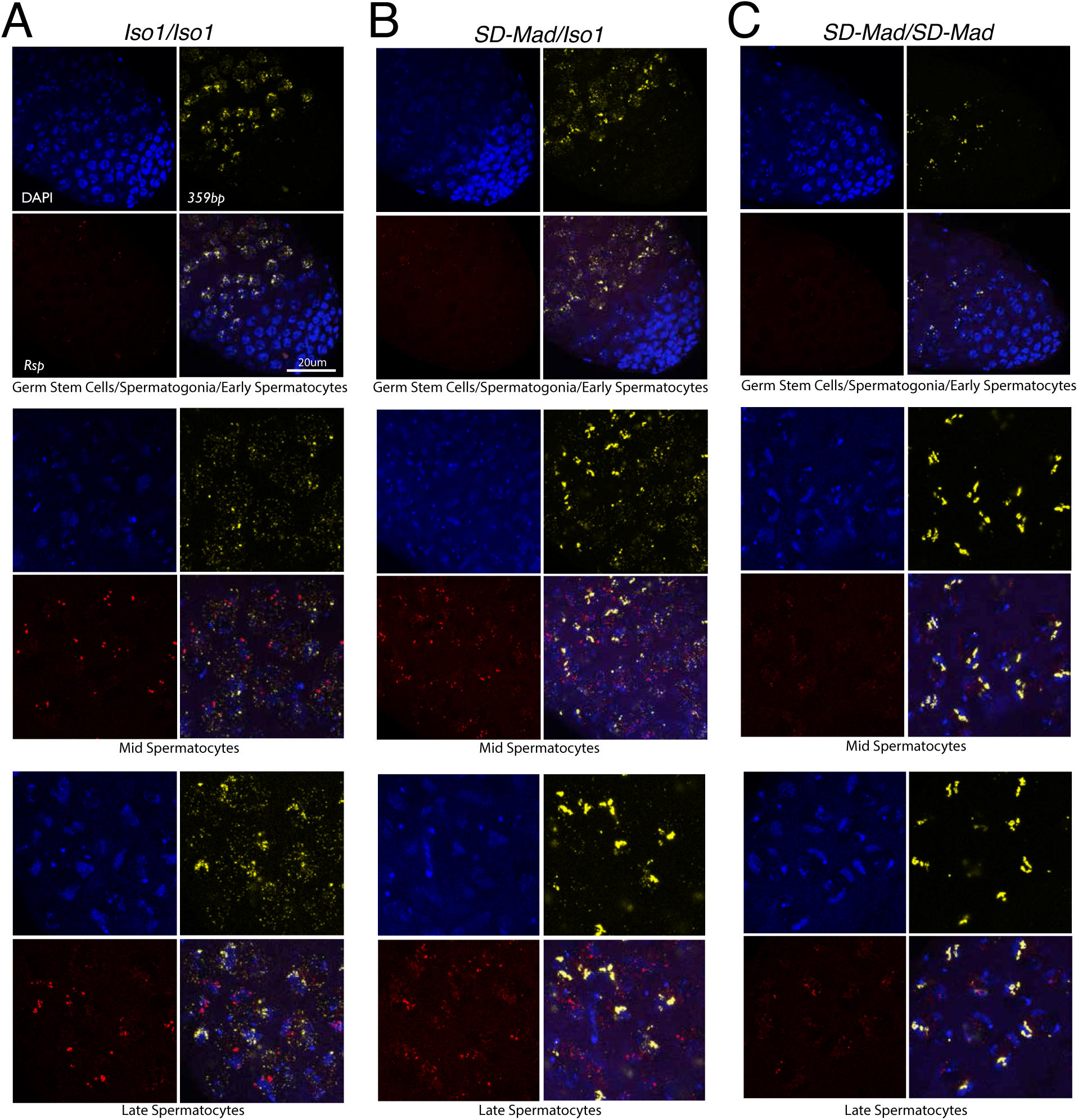
*Rsp* transcripts are not grossly mislocalized in driving backgrounds. DNA FISH in a wild type non driving background (A), a driving heterozygote (B), and a non driving *SD-Mad* homozygote (C). Nuclei are stained with DAPI and fluorescent probes are used to mark *1.C88* (yellow) and *Rsp* (Red). Transcripts are nuclear localized as expected in all stages of expression, from early spermatocytes to late spermatocytes.

**Supplemental Figure 8:**
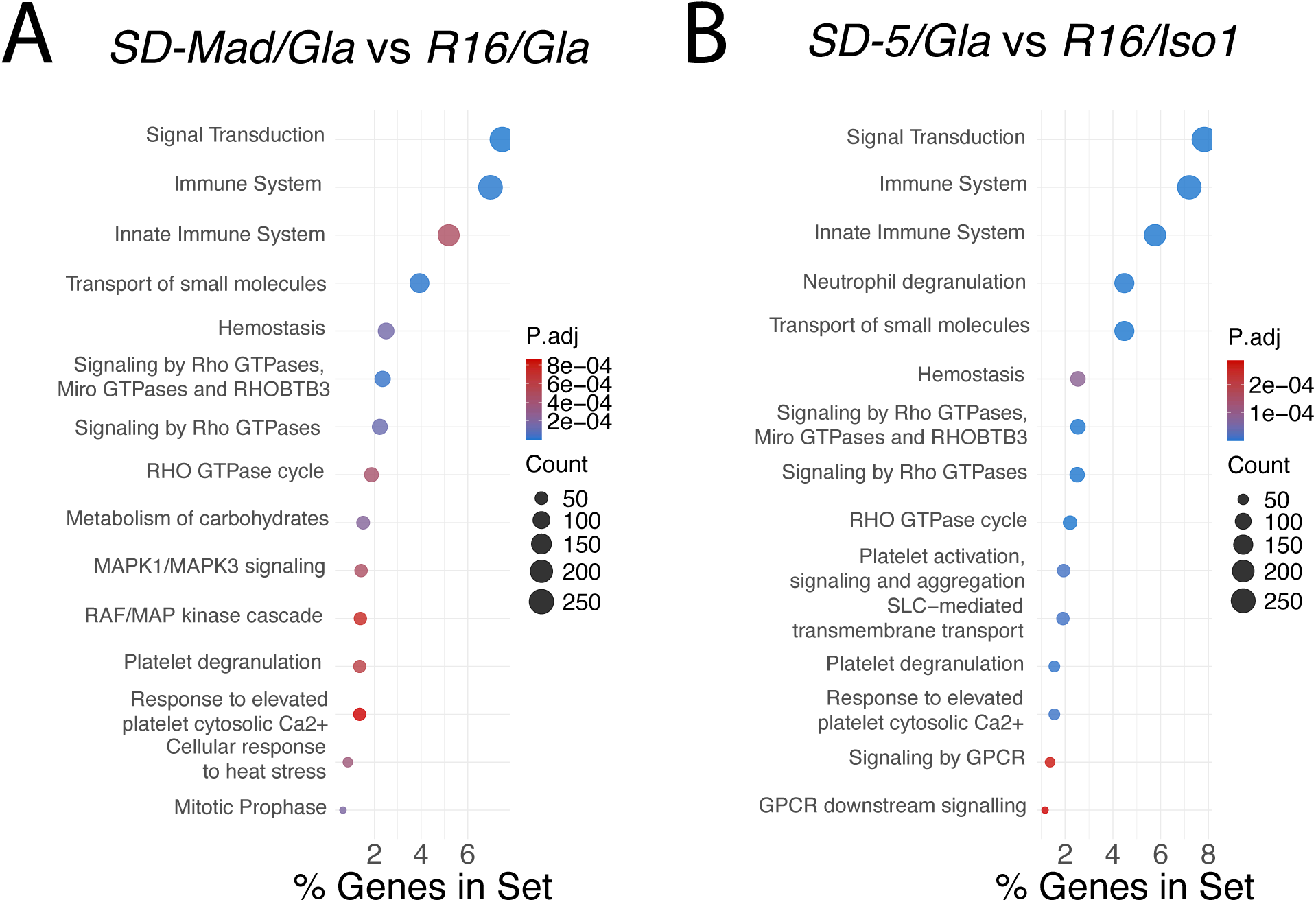
Gene set analysis from the total RNA seq of both *SD/Gla* genotypes shows differences in the immune system and the *Rho GTPase* cycle when compared to *R16/Gla*. A-B) A dotplot of Reactome gene sets shows a broad dysregulation of genes involved in the immune system and *Rho GTPase* cycle. In additon to these common gene sets: *SD-Mad/Gla* also shows a dysregulation in the *MAP* pathway and in mitotic prophase. *SD-5/Gla* shows a dysregulation of G-protein coupled receptor signaling which encompasses a broad set of cellular pathways. The color of the dot represents the *P*-value with blue being smaller *P* values and red being larger *P* values. The size of the dot corresponds to the number of genes that overlap with the given gene set. On the X-axis is the percentage of all differentially expressed genes in the comparison that overlap with the given gene set.

**Supplemental Figure 9:**
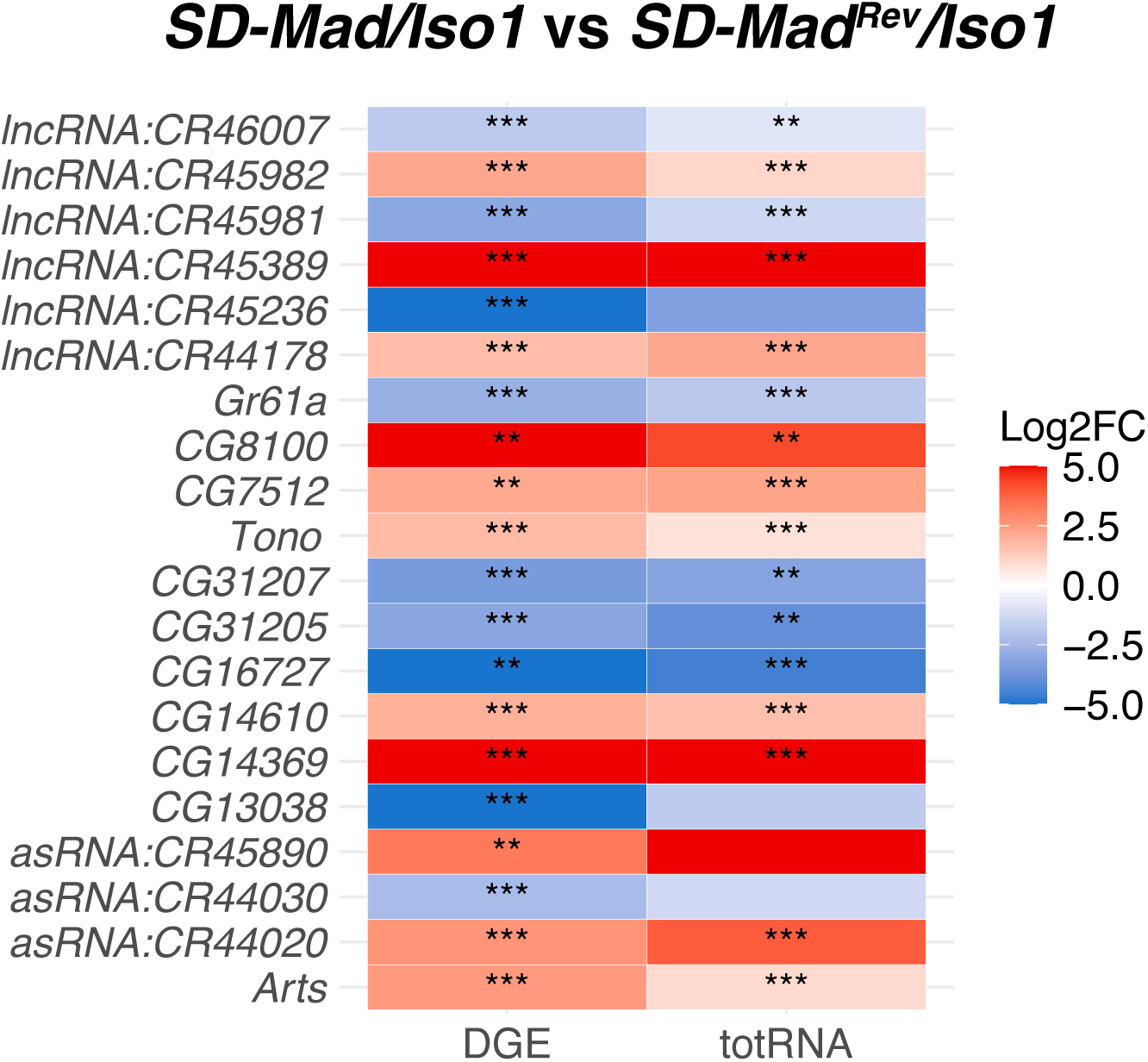
The top 20 most differentially expressed genes are consistent between the digital gene expression and total RNA analysis. A heat plot showing the magnitude and direction of the twenty most differentially expressed genes from the digital gene expression of the SD-Mad/Iso1 vs *SD-Mad^Rev^/Iso1* comparison side by side with the totRNA results. The direction and significance of the differential expression is largely consistent between the totRNA and DGE.

**Supplemental Figure 10:**
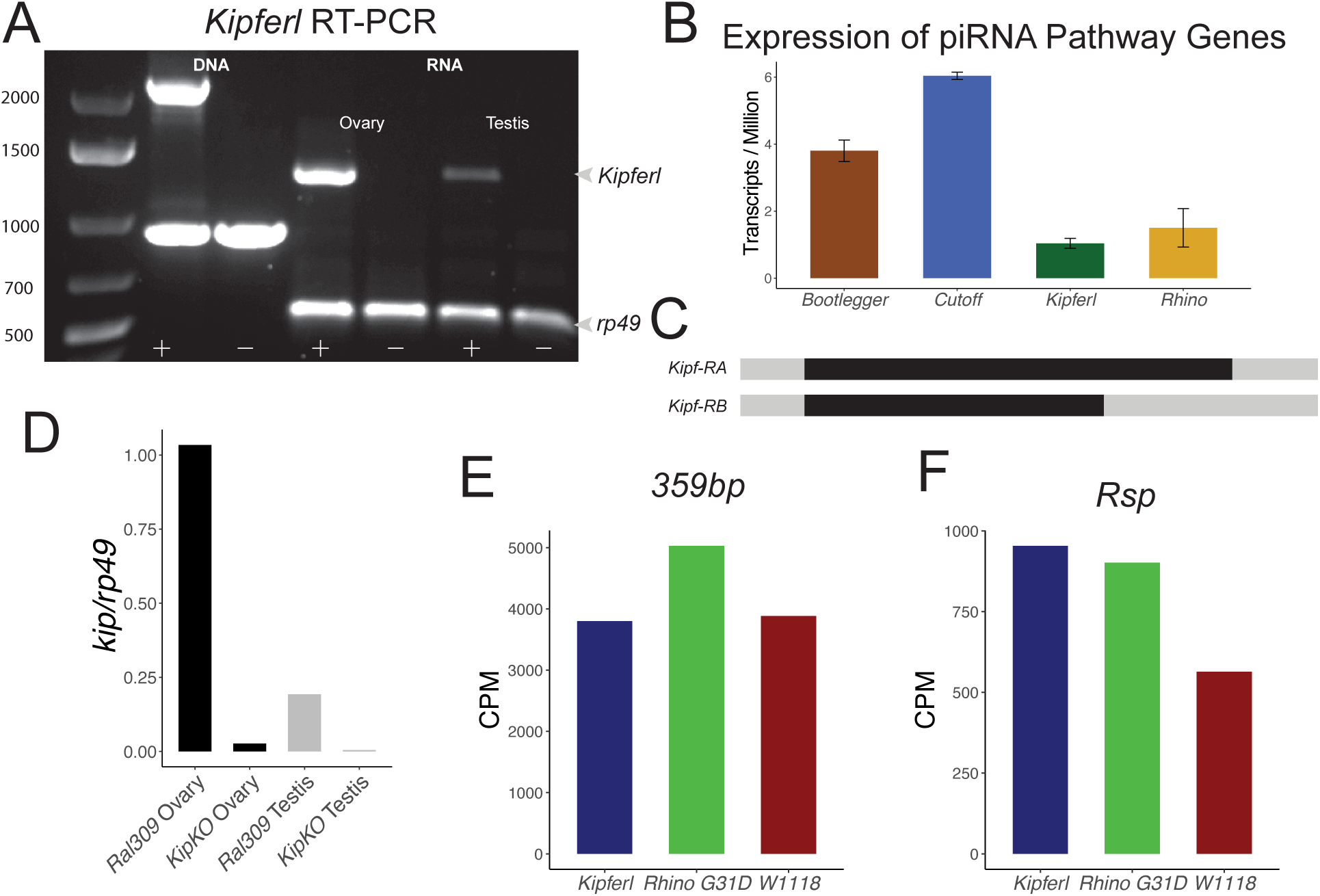
*Kipferl* is expressed in the male germline. A) Genomic PCR for *kipferl* and *rp49* (PCR control) on WT (*Ral3S1*) and *kipf*-KO ovaries and testes. The first two lanes show that the genomic locus is deleted in the knockout. The next two pairs of lanes are RT-PCR reactions from cDNA for ovaries (lane 3-4) and testes (lane 5-6) from the same genotypes. Bands are present in the wild type genotype and absent in the knockout. Quantification of the signal can be found in D. B) Transcripts per million measurements from total RNA seq for various piRNA genes. *Kipferl* is expressed at a slightly lower level than *rhino*. C) Schematics of the two transcripts observed upon amplicon sequencing of the wild type testis *kipf* RT-PCR. They correspond to *kipf*-RA and *kipf*-RB on Flybase. D) Quantification of the nucleotide gel in A. *Kipferl* is expressed at a lower level in testes than in ovaries. E-F) A reanalysis of small RNA seq data from Baumgartner et al. 2024. These single replicate data show an inconsistent affect on the transcription levels of satellite DNA derived small RNAs in a *kipf*-KO background. *359bp* refers to the most abundant component of *1.C88*.

**Supplemental Figure 11:**
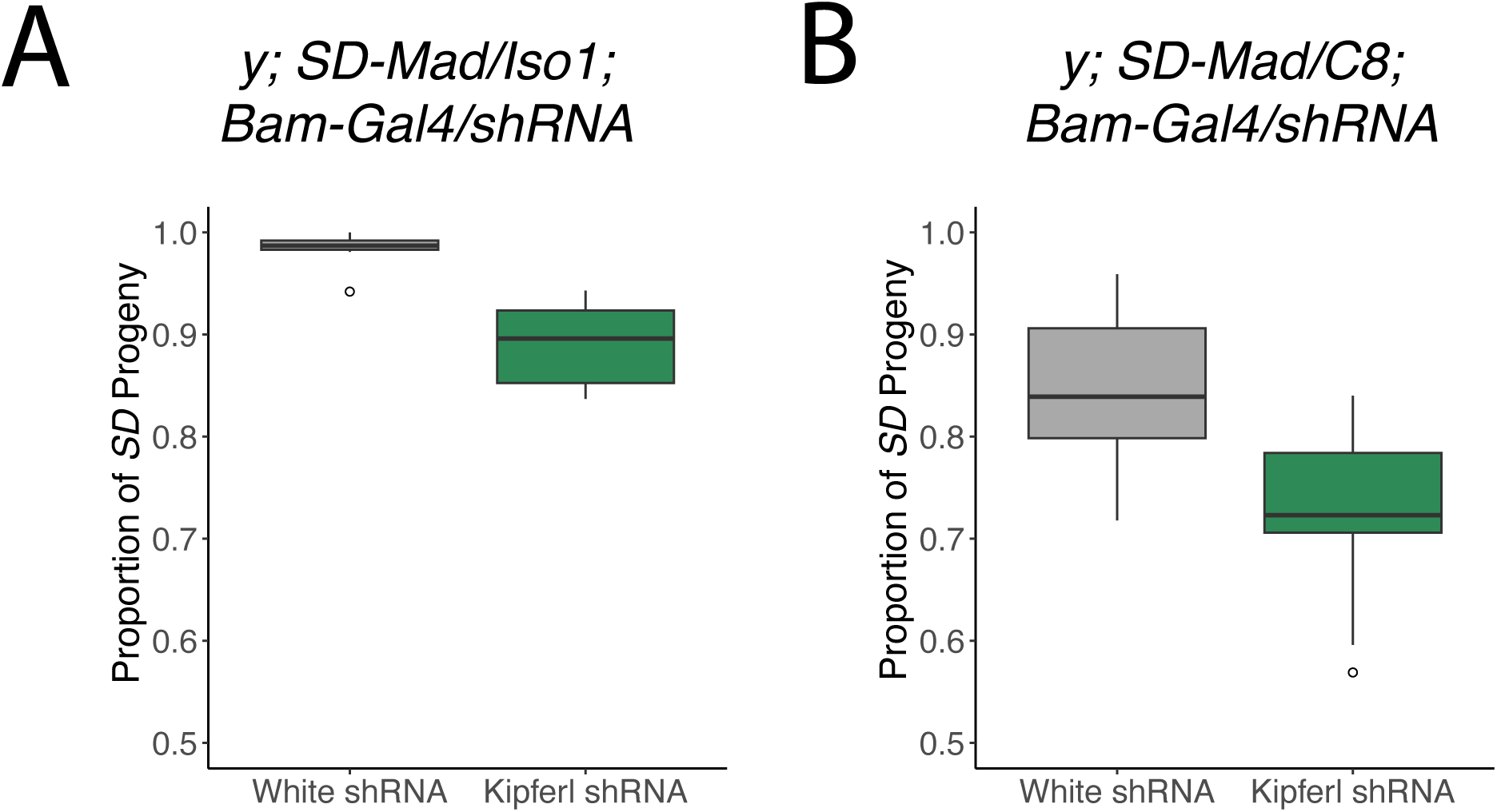
*Kipf*-KD reduces drive strength in both sensitive and semi sensitive backgrounds. A-B) Drive strength analysis for *SD-Mad/Iso1* and *SD-Mad/C8*. *C8* is a *Rsp* deletion line generated from *Iso1* by Eickbush et al and is semi sensitive to *SD*. Short hairpin RNAs for *white* (control) or *kipf* were driven by a *Bam-Gal4/UASp* expression system. Upon *kipf*-KD, *k* value in *SD-Mad/Iso1* drops from *k* = 0.98 to *k* = 0.89 (P < 0.0001 (asin-t)). *K* value in *SD-Mad/C8* drops from *k* = 0.85 to *k* = 0.72 (*P* < 0.0001 (asin-t)).

**Supplemental Figure 12:**
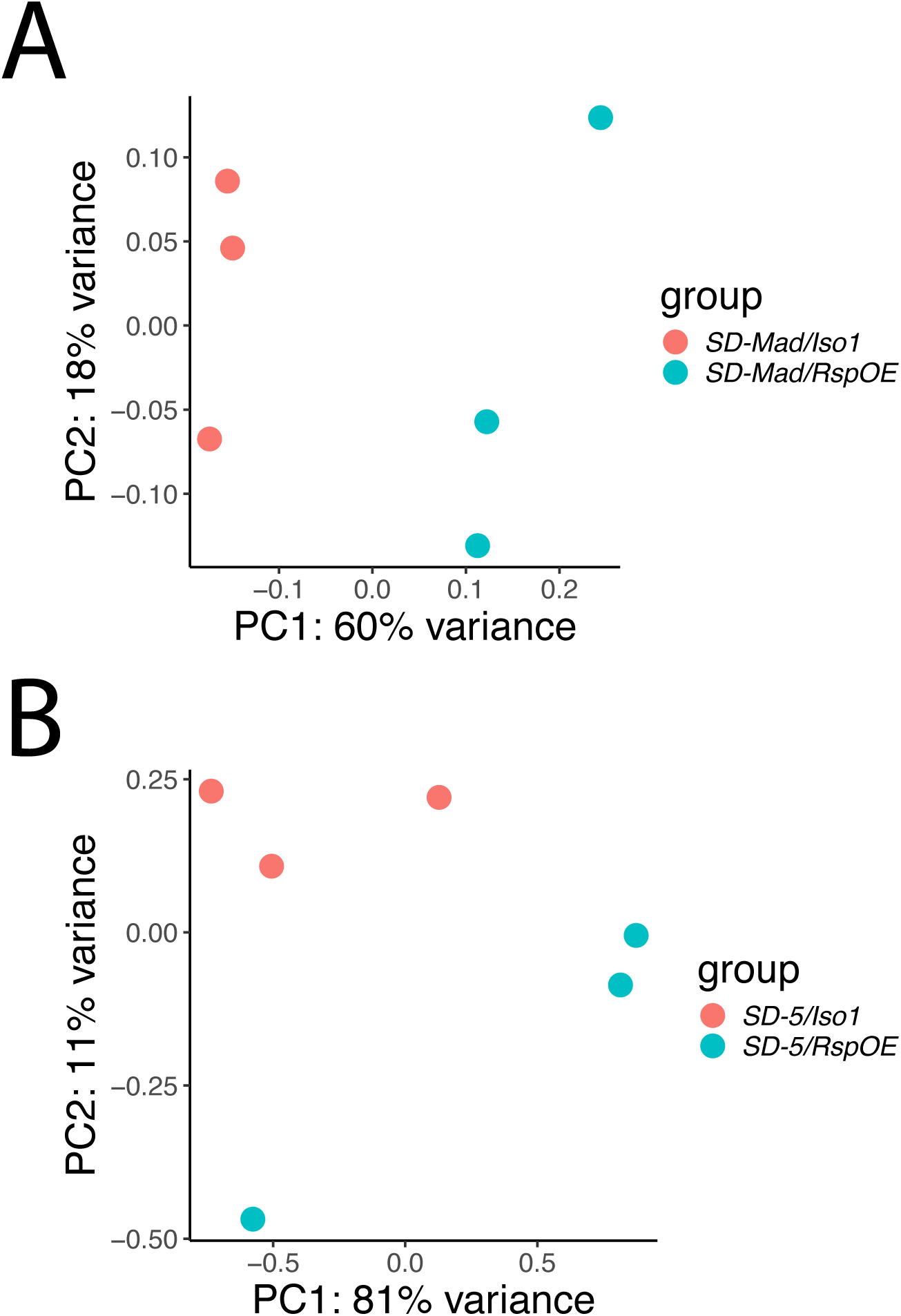
Variance between samples of smRNAs from *SD/Iso1* and *SD/RspOE* backgrounds does not necessarily cluster by genotype. A-B) PCA plots show that the intersample variance between *SD/Iso1* and *SD/RspOE* is largely similar. This is expected as the only difference between these genotypes is a small insertion of *Rsp* into the piRNA cluster, *38C1*.

**Supplemental Figure 13:**
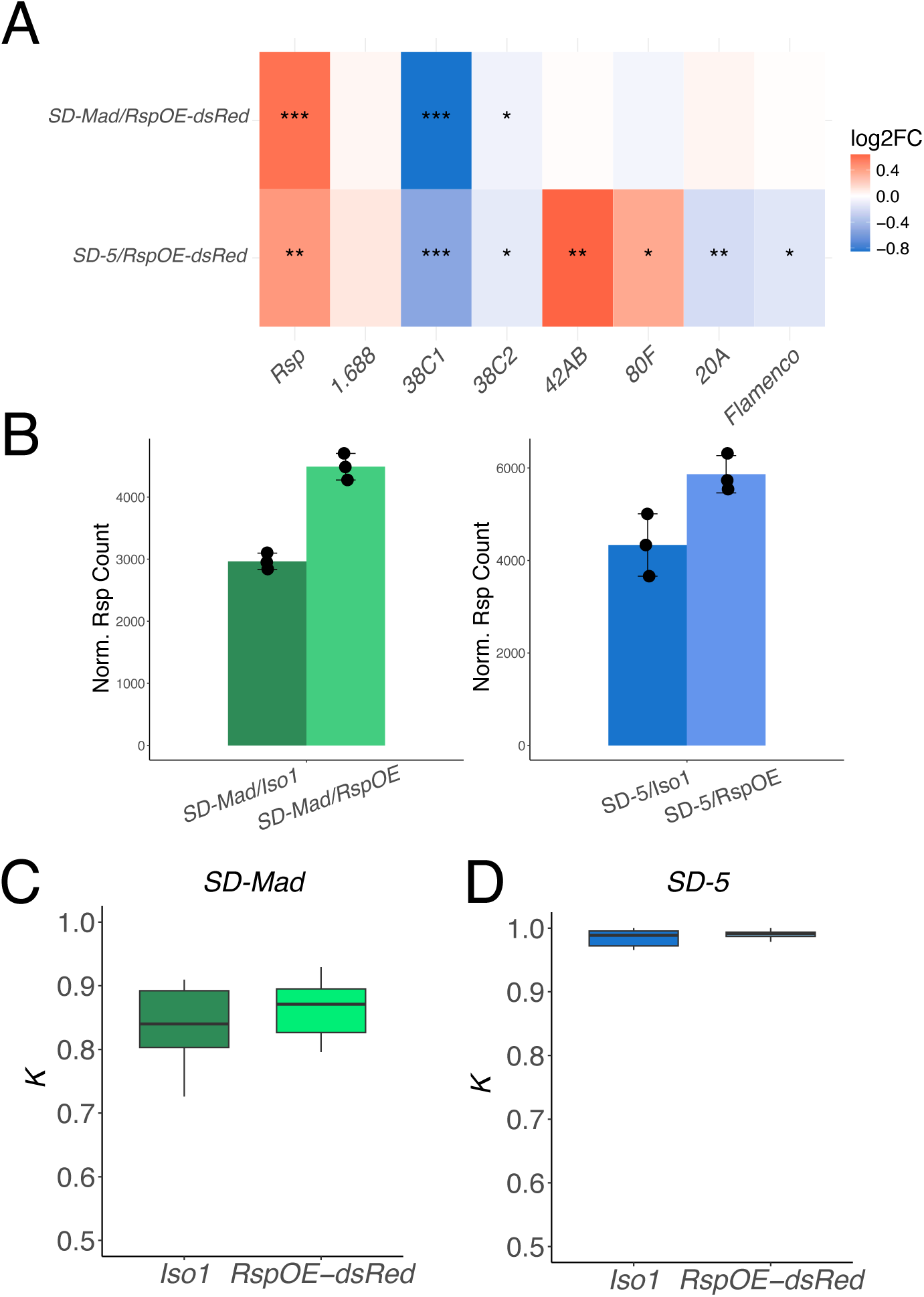
Insertion of both *Rsp* and 3x*P3*-*dsRed* into *38C1* overexpresses *Rsp* but does not lower drive strength. A) A heatplot from small RNA sequencing showing the magnitude and significance of differential expression of satellites and major piRNA clusters between *SD-Mad/Iso1* and *SD-Mad/Iso1^Rsp-dsRed>38C1^* (*RspOE-dsRed*). Similar to the non-*dsRed* results, piRNA cluster *38C1*, and to a smaller degree, *38C2*, are less abundant than expected. B) Bar graphs showing the DESeq2 normalized counts for *Rsp* for *SD/Iso1* and *SD/RspOE-dsRed*. *Rsp* is consistently overexpressed when the 3x*P3-dsRed* marker is present in both *SD* backgrounds. C) Unlike in the experiments where the 3x*P3-dsRed* was removed, drive strength shows no difference between the *Iso1* background (control) and the *RspOE-dsRed* background for either *SD* chromosome. This suggests that something about the 3x*P3-dsRed* insertion abrogates the effect of the *Rsp* overexpression on drive.

